# Using Artificial Intelligence to Assess Cross-Species Transmission Potential of Influenza A Virus

**DOI:** 10.1101/2025.06.17.660059

**Authors:** Jiaying Yang, Pan Fang, Jianqiang Liang, Yihao Chen, Lei Yang, Wenfei Zhu, Mang Shi, Xiangjun Du, Juan Pu, Dayan Wang, Guirong Xue, Zhaorong Li, Yuelong Shu

## Abstract

Influenza A viruses (IAVs) pose pandemic threats through cross-species transmission, yet predicting their adaptive evolution remains challenging. We introduced Influenza A virus Adaptability to host X (FluAdaX), a deep learning framework that integrates a moving average-equipped gated attention mechanism using full-genome sequences. FluAdaX demonstrated robust host classification performance across endemic IAV strains, and outperformed traditional models in detecting avian-to-human transmission. Spillover score and adaptability score were introduced to evaluate host shift risk, which prioritized variants with elevated human adaptation potential, such as H7N9, H9N2 avian IAVs, and H1N1 swine IAVs. Besides HA and NA genes, PB2 and NS genes were found critical for zoonosis. Potential molecular markers associated with avian/human tropism were identified across PB2 and NS genes using XGBoost. FluAdaX provided a dynamic framework to decode IAV host adaptation, enabling real-time risk assessment of cross-species transmission of emerging IAV variants.

## Background

Influenza A viruses (IAVs), single-stranded negative-sense RNA viruses with 8 gene segments encoding over 10 proteins, pose a persistent global public health threat due to their high mutability and cross-species transmission potential^1^. Wild waterfowls are traditionally considered as the primary reservoir for IAVs, which frequently spill over into mammals, leading to epidemics in pigs, dogs, horses^1^, and even pandemics in humans^2–5^, yet we have limited understanding of the underlying mechanisms for host shifts and mammalian adaptation. While sporadic human infections with avian influenza subtypes have been reported, such as H5, H7 and H9^6–10^. The public is concerned about whether these viruses will cause a pandemic, and/or how close they are to a pandemic virus.

The World Health Organization (WHO) developed the Tool for Influenza Pandemic Risk Assessment (TIPRA)^11^ to provide a standardized and transparent approach for assessing the pandemic potential of influenza viruses. However, predicting pandemic potential remains challenging due to the complex mechanism of host adaptation^12–15^. Cross-species transmission, triggered by dynamic mutations and reassortment in the IAV genome, can lead to stable new-species adaptations driven by strong new-host selection factors^5^. Despite progress in identifying adaptive markers^16–19^, systematically detecting such mutations in the increasing population of IAVs is time-consuming and difficult.

Machine learning (ML) methods have emerged as transformative tools for decoding the genomic language of viruses. While conventional ML methods have proven effective in predicting IAV hosts ^20–22^ and identifying key mutations of IAVs to infect humans^23^ based on nucleotide or protein sequences, their reliance on well-studied features limits their capacity to uncover novel biological insights from raw genome sequences. Hie et al. demonstrated the capacity of large language models (LLMs) to predict immune escape patterns of IAV using hemagglutinin (HA) sequence alone^24^. However, a critical bottleneck persisted that the computational inefficiency of standard transformers in handling long sequences, compounded by their weak inductive biases for capturing localized evolutionary constraints^25^. Furthermore, all of these models were trained on single segment (e.g., HA or PB2), ignoring the host adaptation feature in the global context across the whole genome. Considering the frequent mutation and extensive reassortment that complicated the evolution of IAVs, a comprehensive assessment of the whole genome is imperative.

Here, we present FluAdaX, a nucleotide-based deep learning framework that predicts the host adaptation of IAVs originating from human, swine, avian, canine, and equine. Addressing the gap of IAV full-genome modeling, FluAdaX adopts the mechanism of moving average equipped gated attention (MEGA)^26^, which enables efficient capture of both local nucleotide dependencies and global genomic context. This design facilitates the risk assessment of IAV’s host shifts and adaptation through complete genome at nucleotide level. The same framework was also applied to each IAV segment. Critical molecular markers were pinpointed through gradient-based attribution analysis. Our study establishes a unified framework to discern and quantify the host adaptation of IAVs, providing valuable insights into the mechanisms that underlie IAV’s ability to infect and adapt to diverse host species.

## Results

### Overview of FluAdaX

FluAdaX is tailored to learn and evaluate the adaptability of IAV across five common hosts, including human, avian, swine, canine, and equine (Fig. 1). The base module of FluAdaX is a transformer-style model with multiple attention layers. Notably, the architecture of MEGA was employed^26^ as the backbone of FluAdaX. MEGA is a single-head gated attention mechanism equipped with exponential moving average (EMA), which incorporates inductive bias of position-aware local dependencies into the position-agnostic attention mechanism. It can efficiently process extremely long sequences, in terms of computational and information extraction performance. This design enabled FluAdaX to effectively process IAV full-genome sequences, capturing complex patterns of viral adaptation across various hosts (Fig. 1). The FluAdaX outputs were processed with a softmax function to generate a set of probability values (confidence level β) corresponding to the five hosts (Fig. 1a).

**Fig. 1.**
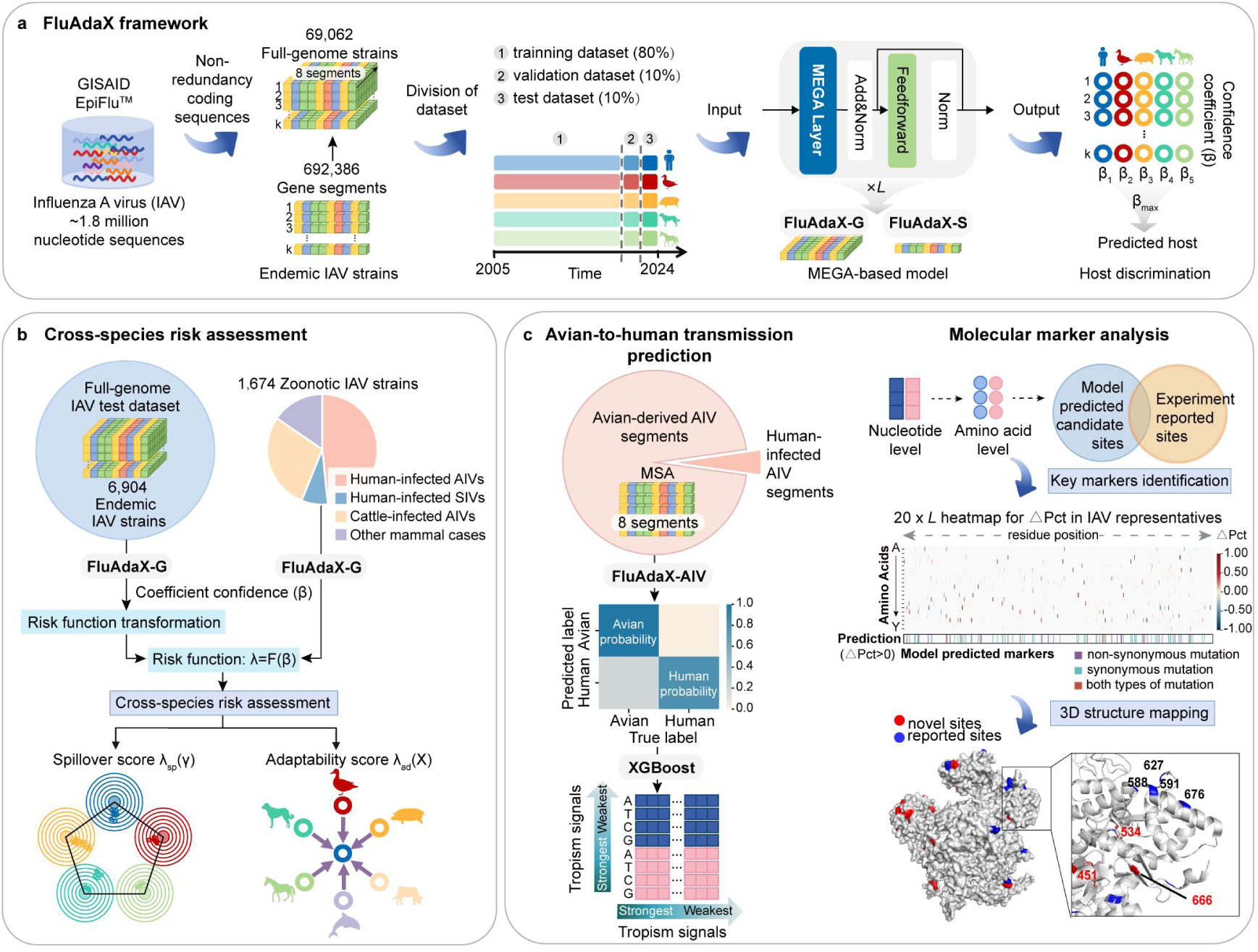
**FluAdaX predicts host adaptability and related molecular markers of influenza A virus based on nucleotide sequences.** a. Workflow of the proposed FluAdaX framework. Endemic influenza A virus (IAV) dataset from five hosts was collected from the GIASID Epiflu database. After qualification, the non-redundancy coding nucleotide sequences were then divided into train dataset, valid dataset, and test dataset according to the collection date chronologically within each host. The backbone of FluAdaX is a transformer-based model incorporating Moving Average Equipped Gated Attention (MEGA) layers, where *L* denotes the number of MEGA layers. FluAdaX-G used a concatenation of alignment-free nucleotide sequences of eight IAV segments as its input. FluAdaX-S used nucleotide sequences of individual segments as its input. FluAdaX-G employed 2 MEGA layers, while FluAdaX-S used 4 MEGA layers. These model were used to predict the adaptability of IAV to different hosts. b. Risk assessment of cross-species transmission of IAVs using FluAdaX-G. Host adaptability of whole-genome strains was converted into risk index to assess the potential of cross-species transmission for epidemic IAV strains and validated using cross-species cases. Spillover score λ_sp_(γ) refers to the spillover potential from epidemic hosts while adaptability score λ_ad_(X) refers to the adaptation potential to target hosts. c. Key molecular markers identification using FluAdaX-AIV model. The model was trained on multiple-sequence alignments (MSAs) of full-length sequences from avian influenza viruses (AIVs) and human infected AIV cases to predict host tropism by gene segment. Gene segments with high human recall were selected to identify host tropism signals at nucleotide level, which were then transformed into amino acid levels. The predicted sites were compared with experimental reported sites and key molecular markers analysis were conducted on the prediction results. Heatmap visualizes ΔPct values across 20×*L* AIV gene segment (where *L* is the protein length). The colored band beneath the heatmap marks potential key sites predicted by the model to with ΔPct>0. Top 15 predicted amino acid sites with ΔPct>0 were then visualized on 3D protein structure.

To develop FluAdaX, endemic IAV strains derived from the selected hosts were collected from EpiFlu Database on Global Initiative on Sharing All Influenza Data (GISAID). After stringent quality control, a total of 692,386 non-redundant coding nucleotide sequences spanning eight IAV genomic segments were curated. Within this dataset, 69,062 endemic IAV strains were identified as full genomes. As illustrated in Fig. 1a, the whole dataset was partitioned into training, validation, and test sets at an 8:1:1 ratio according to the collection timeline within each host category (Fig. 1a; Supplementary Fig. 1b). This chronological partitioning strategy ensured that the training set comprised earlier strains, whereas the validation and test sets included more recent strains. This approach enabled rigorous evaluation on the model whether it could effectively predict the hosts of future emerging variants of IAVs.

FluAdaX-genome (FluAdaX-G) used a concatenation of alignment-free nucleotide sequences of eight IAV segments (∼13kb for each strain) as its input, ensuring that all genetic information and interactions relevant to host tropism were considered. Meanwhile, FluAdaX-segment (FluAdaX-S) model was developed using individual segments as inputs (Supplementary Fig. 1b). The two models were initially tested on the newly collected IAV strains for internal validation. To comprehensively evaluate the performance and generalizability of these models, an external validation was conducted using a dataset comprising zoonotic IAV genomes (Supplementary Fig. 1).

### FluAdaX can accurately identify host taxa of IAVs

FluAdaX-G exhibited robust performance in discriminating hosts of IAVs across five major host taxa, achieving area under the receiver operating characteristic curve (AUC) values of 0.99–1.00 on the test set (n=6,904 sequences) (Fig. 2a,b). This robust performance persisted even when analyzing individual gene segments using the FluAdaX-S model, with AUC values ranging from 0.98 to 1.00 across all segments, indicating the presence of host adaptation signals across the entire IAV genome (Supplementary Table 1). Additional metrics (e.g., recall and precision) further demonstrated classification accuracy. To benchmark FluAdaX-G against other ML approaches, we conducted a comparative analysis using identical training and evaluation datasets across four diverse algorithms, namely, multi-layer perceptron (MLP), Gradient Boosting Machine (GBM), Decision Tree-based Adaptive Boosting (AdaBoost), and Support Vector Machine (SVM). All models achieved comparable performance for host discrimination on endemic IAV strains (Supplementary Table 3). Given that zoonotic spillover events pose significant threat to global public health, we specifically evaluated the capacity of FluAdaX to identify cross-species transmission events of IAVs despite its original design focused on host adaptation. An external validation set comprising 940 zoonotic IAV strains including human-infected avian influenza viruses (AIVs) and human-infected swine influenza viruses (SIVs) was constructed for this evaluation (Fig. 2a). For avian-to-human transmission cases, FluAdaX-G achieved an overall recall of 0.73. In swine-to-human transmission scenarios, the model achieved a moderate overall recall of 0.23. FluAdaX-S model and classical ML methods exhibited substantially reduced prediction capacity for avian-to-human transmission compared to full-genome prediction while maintaining competitive performance in swine-to-human transmission scenarios (Fig. 2c; Supplementary Table 2).

**Fig. 2.**
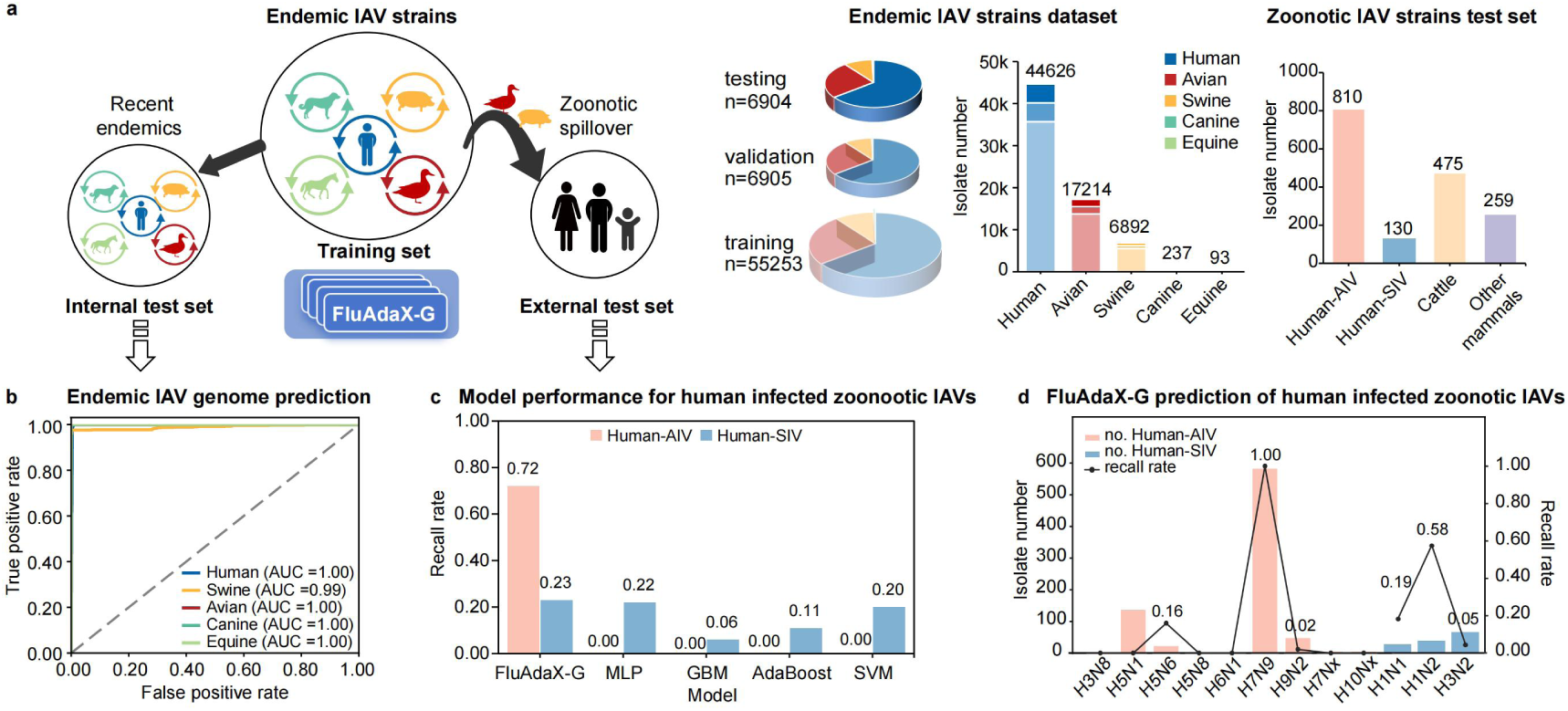
**Host prediction results of FluAdaX-G.** a. Full-genome influenza A virus (IAV) datasets for FluAdaX-G. A total of 69,062 endemic IAV strains were leveraging for training, validating, and testing FluAdaX-G, while an additional dataset of 940 human-infected zoonotic strains was tested for external validation. Pie charts and the adjacent bar graph show the composition of training, validation, and testing set of FluAdaX-G. The external test set comprising zoonotic IAV strains are shown on the right. b. Receiver Operating Characteristic (ROC) curves of FluAdaX-G based on the full-genome sequences of IAVs. Each ROC curve reflects the model’s prediction capacity for a specific host, with individual AUC values quantifying the performance for each host source of IAVs. The higher AUC values indicates better prediction power. c. Performance of various machine learning models in identifying cross-species transmission cases from animals to humans, where a higher recall rate signifies a stronger abilities to detect human-infected cases. The pink bars represent the recall rates for human-infected avian influenza viruses (Human-AIV). The blue bars represent the recall rates for human-infected swine influenza viruses (Human-SIV). d. Host recall rates of FluAdaX-G for cross-species transmission in humans by different subtypes of avian and swine influenza viruses. The bar graphs show the number of tests carried out for human infection with avian influenza viruses (pink), and swine influenza viruses (blue).

The model presented distinct performance patterns across subtypes. For human-infected AIVs, the highest recall rate (1.00) was observed for H7N9, followed by H5N6 (0.16) and H9N2 (0.12). Human infections with other AIV subtypes were rarely identified by the model (Fig. 2d). We presumed that human-infected H7N9 AIVs contained conserved mammalian-adaptive mutations that generate strong adaptive signals^27^, while the low sensitivity for most AIV subtypes may stem from their frequent reassortment with wild bird lineage^1^, obscuring host-specific signals.

For human-infected SIVs, H1N2 (recall=0.58) showed the highest recall rate, followed by H1N1 (recall=0.19) subtype (Fig. 2d). Frequent human-to-swine reverse zoonoses of H1N1 IAVs may also confound the host boundaries for model prediction^28^.

### Evaluating host adaptability of endemic IAVs based on FluAdaX-G

As above, FluAdaX-G exhibited comparable performance in host discrimination on endemic IAV strains and showed superiority in assessing cross-species transmission risks compared to other models. Therefore, we established a quantitative framework to assess cross-species transmission potential of each IAV strain based on this model (Fig. 1b). We established a dual-metric framework disentangled two critical dimensions of cross-species transmission: spillover and adaptation potential. Two novel metrics, namely, spillover score λ_sp_(γ) and adaptability score λ_ad_(X), were calculated (Methods). Successful cross-species transmission and consequent adaptation are rare but disastrous, therefore we calibrated the outputs of FluAdaX-G to prioritize sensitivity in detecting these signals. The adaptation level within each selected host was referenced to average confidence level, which serves as a validated reference standard because the average confidence level to native hosts was the highest when applied to the internal test set of 6,904 endemic IAV strains (Supplementary Fig. 2a).

The spillover score λ_sp_(γ) quantified the evolutionary constraint of an IAV strain within its native host γ, by measuring its fitness deficits compared to host γ-adapted consensus variants. This metric was normalized to a range of 0-10 across our 6,904 endemic IAV test sets, with a higher value indicating a greater risk of spillover (i.e., reduced adaptation to the native host γ). Conversely, the adaptability score λ_ad_(X) measured the relative risk of IAV’s capacity to overcome host-specific barriers and establish adaptation to a target host X, rather than causing transient infection. The λ_ad_(X) was also normalized to the interval [0,10] by leveraging the endemic IAV strains, with a higher value reflecting a higher potential of adaptation to host X.

After normalization, endemic IAV strains across five hosts shared an identical mean λ_sp_(γ) value (0.428), whereas the 95% confidence intervals (CIs) varied. Human strains exhibited the narrowest 95%CI (0.428-0.428). Avian strains exhibited minimal variation (95%CI: 0.427-0.429), while canine and equine strains both had 95%CI of 0.425-0.431. Swine strains, however, showed greater variability with a wider 95%CI of 0.297-0.559 (Supplementary Table 4). The adaptability score strongly correlated with host adaptation across endemic IAV strains, as evidenced by the highest mean λ_ad_(X) value to their native host. For example, human endemic IAVs had a mean λ_ad_(Human) value of 7.607 (95%CI: 7.606,7.607), whereas swine endemic IAVs showed a mean λ_ad_(Human) value of 0.653 (95%CI: 0.481,0.825). Avian, canine, and equine endemic IAVs exhibited lower mean λ_ad_(Human) values (Supplementary Table 4; Supplementary Fig. 2b). In this study, an IAV strain isolated from host γ is considered to pose elevated spillover risk to a target host if its spillover score exceeds the upper 95% CI bound of endemic IAVs circulating in host γ. Meanwhile, the strain exhibits elevated adaptation potential if its λ_ad_(X) score surpasses the upper 95% CI bound of λ_ad_(X) values in endemic strains from host γ. A few swine endemic IAVs showed elevated adaptability potential to other hosts, consistent with their role as cross-species intermediates^29^. Among them, H1N1 SIVs emerged as representatives of this adaptive plasticity, exhibiting significantly higher λ_ad_(Human) and higher λ_sp_(swine) than other SIV subtypes (Fig. 3b; Supplementary Fig. 3).

**Fig. 3.**
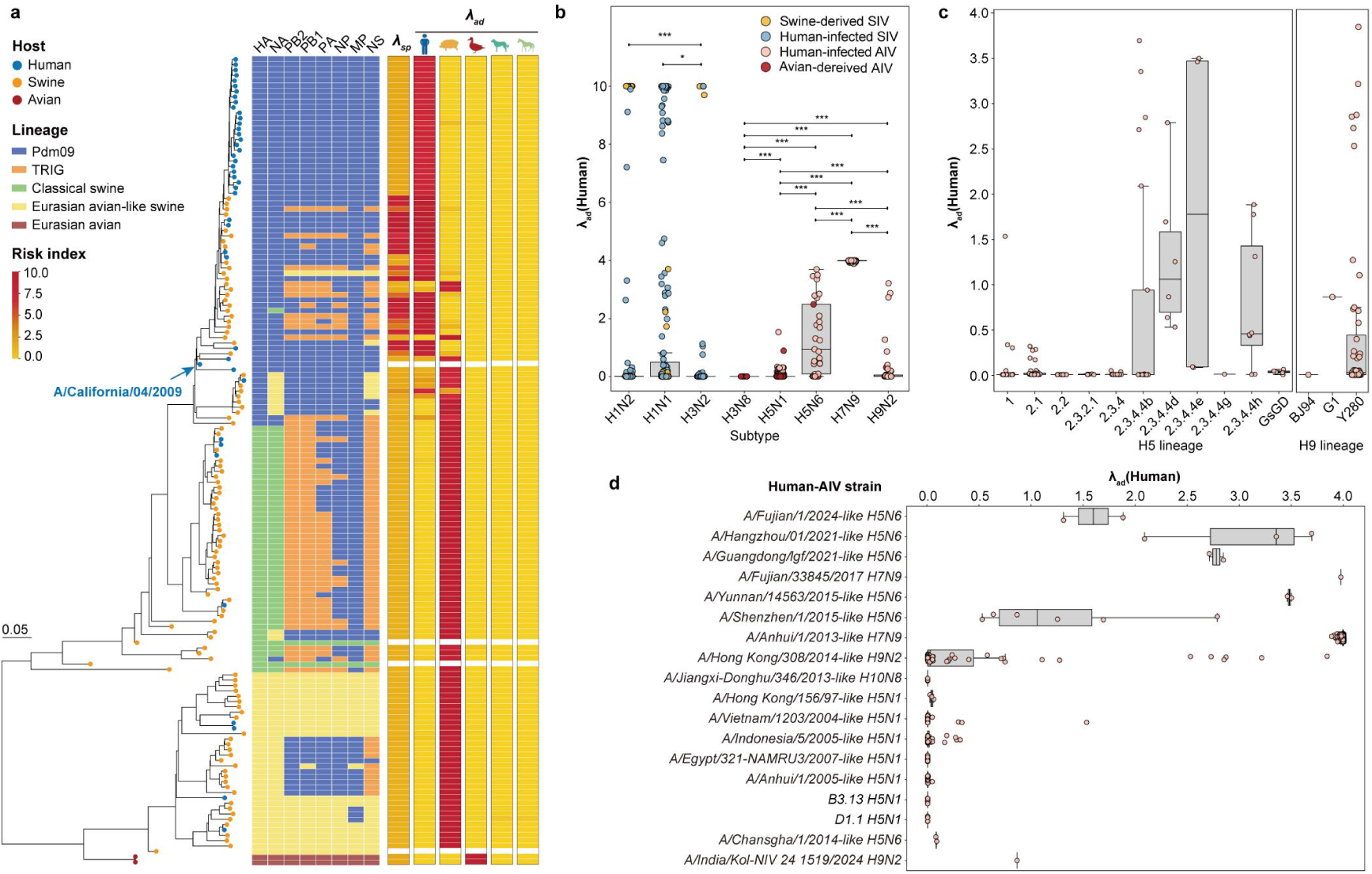
**Risk assessment of influenza A Viruses (IAVs) adaptation to humans.** a. Phylogeny of whole genome sequences of H1N1 subtype IAVs (n=151) and the corresponding risk scores evaluated by FluAdaX-G. Maximum likelihood tree of HA genes of H1N1 viruses were shown on the left, encompassing 147 test H1N1 IAV strains from human, swine, avian, and an additional 4 reference strains used as a backbone. Tip points are colored by hosts. Various lineages of each segment are represented by different colored tiles adjacent to the tree. The six gradient columns on the right represent the spillover score λ_sp_(γ) from the original host and adaptability scores λ_ad_(X) towards five target hosts (human, swine, avian, canine, equine) as evaluated by FluAdaX-G. For λ_sp_(γ), yellow indicates low spillover risk from (/high adaptability to) orignal host, and red means the opposite. For λ_ad_(X), yellow indicating low adaptability and red indicating high adaptability to the target host. The phylogenetic tree of all segments with more complete information are shown in Supplementary Fig. 4-11. b. Comparative adaptation risk of IAV subtypes to humans. The boxplot and scatter plot jointly illustrate the distribution of adaptation risks across different IAV subtypes. The colors of the scatter points represent the origin of the strains, including avian-derived AIV, human-infected AIV, swine-derived SIV, or human-infected SIVs. The upper and lower whiskers of the boxplot represent the upper and lower quartiles, respectively, with the line inside indicating the median. Differences between subtypes were compared using Mann-Whitney U tests, with * indicating *P*<0.05, ** indicating *P*<0.01, and *** indicating *P*<0.001. c. Human adaptation risk of human-infected AIVs across different HA lineages. The boxplot and scatter plots on the left represent the risk among H5 lineages, while thoses on the right represent the risk among H9 lineages. The upper and lower whiskers of the boxplot represent the upper and lower quartiles, respectively, with the line inside indicating the median. d. Human adaptation risk assessment of major human-infected AIV strains. Strain names of representatives were shown. The boxplot and scatter plots on the right represent the human adaptability score λ_ad_(X) corresponding to those strains. The upper and lower whiskers of the boxplot represent the upper and lower quartiles, respectively, with the line inside indicating the median. Risk assessment of all human-infected AIV strains with more complete information are shown in Supplementary Fig. 13.

While these metrics were utilized for risk assessment of cross-species transmission, several considerations should be noted. The model prioritized variants that exhibit concurrent elevation of spillover score and adaptability score, as exemplified by H1N1 SIVs. However, these scores inferred non-directional interactions among hosts, which should be carefully interpreted. Some endemic IAVs may retain ancestral host adaptations while gaining cross-species transmission capabilities, leading to potential mismatches between predicted adaptability scores and observed transmission outcomes. Integration with spatiotemporal circulation patterns and functional validation remains essential for real-world risk assessment.

### Reverse zoonotic transmission drives adaptive plasticity in SIVs

To dissect the evolutionary mechanisms underlying the elevated λ_ad_(Human) of H1N1 SIVs, we conducted phylogenetic analysis of H1N1 IAV genomes (Fig. 3a; Supplementary Fig. 4-11). Some pdm09 lineage H1N1 subtype SIVs exhibited high adaptation potential to human with λ_ad_(Human) value ranging from 7.45 to 10.00, alongside elevated spillover risk from swine [λ_sp_(Swine)=1.95-9.88], whereas other H1N1 SIV lineages, such as classical swine and Eurasian avian-like swine lineages, remained low adaptability to human (Fig. 3a). These human-adapted SIV strains contained nearly all external genes originated from human H1N1pdm09 viruses, incorporating with diverse internal gene constellations from pdm09, Eurasian avian-like swine, and triple reassortant internal gene (TRIG) lineages. This phenomenon could be explained by their likely origin from reverse zoonotic transmission of human pdm09 strains into swine, retaining ancestral host adaptation while selective acquisition of critical mutations facilitated adaptation to novel hosts. Critically, swine-to-human transmission events validated the adaptive plasticity of these high risk strains, with median values of λ_sp_(Swine)=6.29 (IQR: 1.01-9.80), λ_ad_(Human)=9.37 (IQR: 3.86-10.00), and λ_ad_(Swine)=0.63 (IQR: 0.01-6.14) providing quantitative support. We supposed that this genomic pattern enabled bidirectional fitness optimization at the human-swine interface. Some pdm09 lineage SIV have developed high adaptation to swine, which might be driven by critical mutation combinations across viral genomes and recurrent reassortment, rather than uniform segment contributions or phylogenetic proximity alone.

### FluAdaX-G quantifies adaptation dynamics in zoonotic IAVs

We compared the human adaptability score λ_ad_ (Human) between human-infected strains and their counterparts circulating in animal populations to evaluate the human adaptability of these zoonotic viruses following cross-species transmission. Our findings revealed increased human adaptation in both avian and swine IAVs isolated from human cases. Following spillover to humans, avian strains showed increased λ_ad_(Human) from 0.011 (95%CI: 0.007, 0.014) to 2.966 (95%CI: 2.849, 3.084), while swine strains rose from 0.653 (95%CI: 0.481, 0.825) to 2.360 (95%CI: 1.633, 3.087), (Supplementary Table 4). FluAdaX-G exhibited significant concordance in prioritizing high-risk variants with the WHO’s TIPRA framework^11^, particularly H7N9, Y280 lineage H9N2, and clade 2.3.4.4 H5 viruses, where human adaptability scores strongly correlated with TIPRA’s human infection risk index (Pearson’s *r*=0.87, *P*<0.05, Supplementary Table 5). This concordance indicated FluAdaX-G’s capacity to decode host adaptation processes through a genomic lens. Divergence observed in clade 2.3.2.1 H5N1 likely reflect temporal sampling biases that captured distinct phases of adaptive evolution.

Subtype- and lineage-specific host adaptation dynamics among influenza viruses were evident (Fig. 3b-c). Among SIVs, H1N1 (mean λ_ad_(Human)=1.89) and H1N2 (mean λ_ad_(Human)= 0.94) exhibited significantly higher human adaptation potential than H3N2 subtype SIVs (mean λ_ad_(Human)=0.18; *P*<0.05) (Fig. 3b). Human-infected H1N2 subtype SIVs displayed a statistically higher risk of adaptation to human (λ_ad_(Human)=5.80) relative to human-infected H1N1 subtype SIVs (λ_ad_(Human)=2.10, *P*<0.001), with most of the high risk H1N2 SIVs were isolated before 2010 and one of them was collected in 2024 in USA (Supplementary Fig. 12d). For AIVs, H7N9 presented the highest human adaptability (mean λ_ad_(Human)=3.99), followed by H5N6 (1.36) and H9N2 (0.29) (Fig. 3b). Phylogenetic analysis revealed HA lineage-specific adaptation disparities among human-infected H5 strains (Fig. 3c). H5 2.3.4.4 lineage demonstrating significantly elevated human adaptability score compared to other H5 lineages, with clade 2.3.4.4e exhibited the highest human adaptability score (mean λ_ad_(Human)=1.79), followed sequentially by 2.3.4.4d (mean λ_ad_(Human)=1.30) and 2.3.4.4b (mean λ_ad_(Human)=1.12). Some of H9 Y280 lineage strains and a recent H9 G1 variant showed elevated human adaptation potential.

Comparative analysis of classical and emergent zoonotic AIV strains revealed distinct human adaptation potential (Fig. 3d). H7N9 variants including A/Anhui/1/2013-like H7N9 and A/Fujian/33845/2017 H7N9, H5N6 variants such as

A/Yunnan/14563/2015-like, A/Guangdong/lgf/2021-like, and A/Hangzhou/01/2021-like viruses, as well as the newly emergent H9N2 variant A/India/Kol-NIV 24 1519/2024 exhibited markedly elevated human adaptability scores. Other classical strains, such as A/Hong Kong/308/2014-like H9N2 and A/Vietnam/1203/2004-like H5N1, showed moderate human adaptability score. These high risk variants consisted of diverse gene constellations (Supplementary Fig. 13), which obscured the contribution of individual gene segments to cross-species transmission. In contrast, recent H5N1 strains, including the emergent North American B3.13 and D1.1 variants, displayed stable avian adaptation, with model predictions overwhelmingly favouring avian over other mammalian hosts (human/swine/canine/equine). As indicated above, coordinated mutations in specific gene constellations boosted zoonotic potential, whereas H5N1’s ecological persistence in avian hosts constrained its pandemic emergence capacity (Fig. 3d).

### Four of eight segments act as dominant contributors for AIV human adaptation besides external genes

To further analyze the critical gene segments across diverse gene constellations underlying human adaptation of IAVs, FluAdaX-AIV models were developed for each gene segments focusing on the host discrimination in avian-to-human transmission (Fig. 1c). Previous FluAdaX-S model trained solely on endemic IAV strains exhibited limited accuracy in identifying avian-to-human transmission events. To address this, human-infected AIV sequences were added for model training. Multiple sequence alignments of eight independent genes were constructed from 113,722 avian-derived and 5,585 human-infected AIVs (Supplementary Table 6). For the HA and NA segments, H9 and N2 genes were selected as the representatives for modeling and analysis. The FluAdaX framework was subsequently applied to these aligned sequences to identify host within each nucleotide alignment.

Our findings revealed that the contributions of AIV segments to avian-human adaptation may be distinct. For avian-derived AIVs, all segment exhibited strong avian adaptation signals, with recall rates ranging from 0.73 to 1.00 (Fig. 4a). For human-isolated AIVs, HA and NA demonstrated great significance in cross-species transmission to humans, with recall rates both exceeding 0.80. Notably, PB2 and NS genes outperformed other internal genes, achieving recall rates of 0.70 and 0.57, respectively, and were identified as critical contributors of human infection with AIVs (Fig. 4a). We further focused on PB2 and NS genes, and found human prediction disparity across lineages (Fig. 4b,c). Phylogenetic analysis suggested that PB2 and NS genes with elevated human probability (>0.5) predominantly originated from H9N2 G57-like genes, while PB2 gene from a Eurasian wild bird 1-derived cluster showed comparable human probability. Additionally, PB2 genes derived from chicken H9N2 lineage, a minor group from waterfowl H6-derived NS genes, and sporadic genes from other lineages also exhibited elevated human probability. These results further highlighted the dynamic pattern of viral adaptations.

**Fig. 4.**
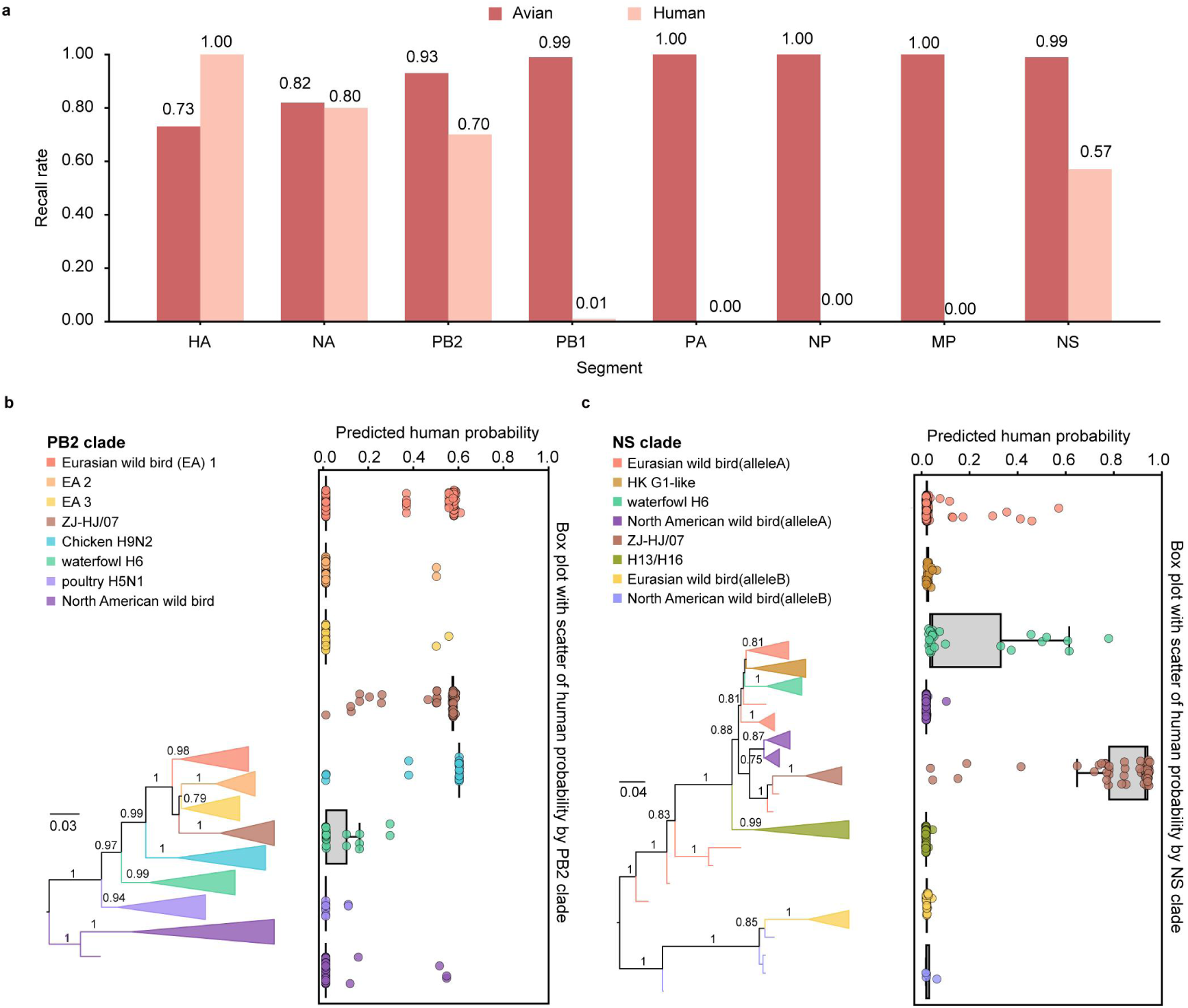
**Model prioritized genes for avian-to-human cross-species transmission of avian influenza viruses.** a. FluAdaX-AIV model performance in segment-specific host prediction for avian and human infections caused by avian influenza viruses. Recall rates quantify model sensitivity in identifying host through individual segment analysis. Avian-derived AIVs are in red, human-infected AIVs are in pink. Detailed information see Supplementary Table 7. b. Human probability prediction correlates of phylogeny. Maximum likelihood phylogeny of PB2 genes (n=2,204) (left) is paired with human probability distribution (right). For clarity, major clades are collapsed and denoted by distinct colors. While boxplots show human probability distributions across major clades (median±IQR), scatter plots map tip-specific human probabilities. c. Human probability prediction correlates of phylogeny. Maximum likelihood phylogeny of NS genes (n=1,746) (left) is paired with human probability distribution (right). For clarity, major clades are collapsed and denoted by distinct colors. While boxplots show human probability distributions across major clades (median±IQR), scatter plots map tip-specific human probabilities.

### Machine learning prioritizes PB2/NS mutations critical for AIV human adaptation

Key mutations will determine virus’s features and enable it to infect new species as the host tropism shifts. Although four segments were predicted to be dominant contributors for AIV human adaptation, we focused on PB2 and NS genes due to their capacity to reassort with multiple AIV subtypes. Avian and human representative sequences were selected via influence function (Methods). Using the eXtreme Gradient Boosting (XGBoost) algorithm, we then explored the key molecular markers related to IAV host tropism between avian and human. Model training yielded non-zero feature importance scores quantifying the contribution of individual molecular marker at nucleotide level, identifying 160, 157, and 60 nucleotide sites on the PB2, NS1, and NS2 genes. Notably, most of these sites localized to the third codon position (Supplementary Fig. 14b; Supplementary Table 9). Subsequent mapping of these nucleotide markers to protein sequences revealed both synonymous and non-synonymous amino acid mutations across functional domains

(Supplementary Fig. 14c-f). These markers were then interpreted into amino acids and ranked by feature importance scores (Methods). Collectively, 155, 128, and 49 candidate amino acid positions were identified in PB2, NS1, and NS2 proteins, respectively, associated with host tropism between avian and human (Fig. 5a; Supplementary Table 8-10). Among the top-ranked predictions, 31/100 sites in the PB2 protein, 20/100 sites in the NS1 protein, and 5/49 sites in the NS2 protein had been experimentally reported to be related to human adaptation. Some residues with dual-host compatibility might be omitted by the model. To delineate human-adapted signatures, we analyzed codon composition divergence at each amino acid site between representative IAV sequences isolated from avian and human. The ΔPct values ranging from -1 to 1 were calculated to describe codon frequenncy bias at each amino acid site, where ΔPct>0 indicated elevated proportion in human-infected AIV sequences compared to avian-derived AIV sequences (Fig. 5b). Heatmap visualization showed that a large proportion of the predicted residues exhibited higher proportion in human (ΔPct>0), with non-synonymous mutations prioritized as potential molecular markers for avian-to-human transmission. Non-synonymous mutations on 29, 39, and 16 residues were identified in the PB2, NS1, and NS2 genes, respectively (Supplementary Table 11). Notably, classical human-adaptive genetic markers in the PB2 gene (e.g., E627K, D701N, K588R, T473K) ranked within the top four prediction (Supplementary Table 11), aligned with their significance roles in mammalian adaptation across experimental^18,19,30^. Other candidate molecular markers in PB2 and NS genes need further experimental verification. The top 15 ranked residues of each selected gene were visualized on the 3D protein structure (Fig. 5c). Structural mapping revealed a dispersed distribution across functional domains rather than localized clustering for PB2 genes, despite more experimental verification had been conducted on the mutations located in the 627 domain and cap-binding domain. 50% (24/48) of the candidate molecular markers in the NS1 protein localized to the effector domain, majority (12/18) of the NS2 predictions while localized to the N-terminal (Supplementary Table 11).

**Fig. 5.**
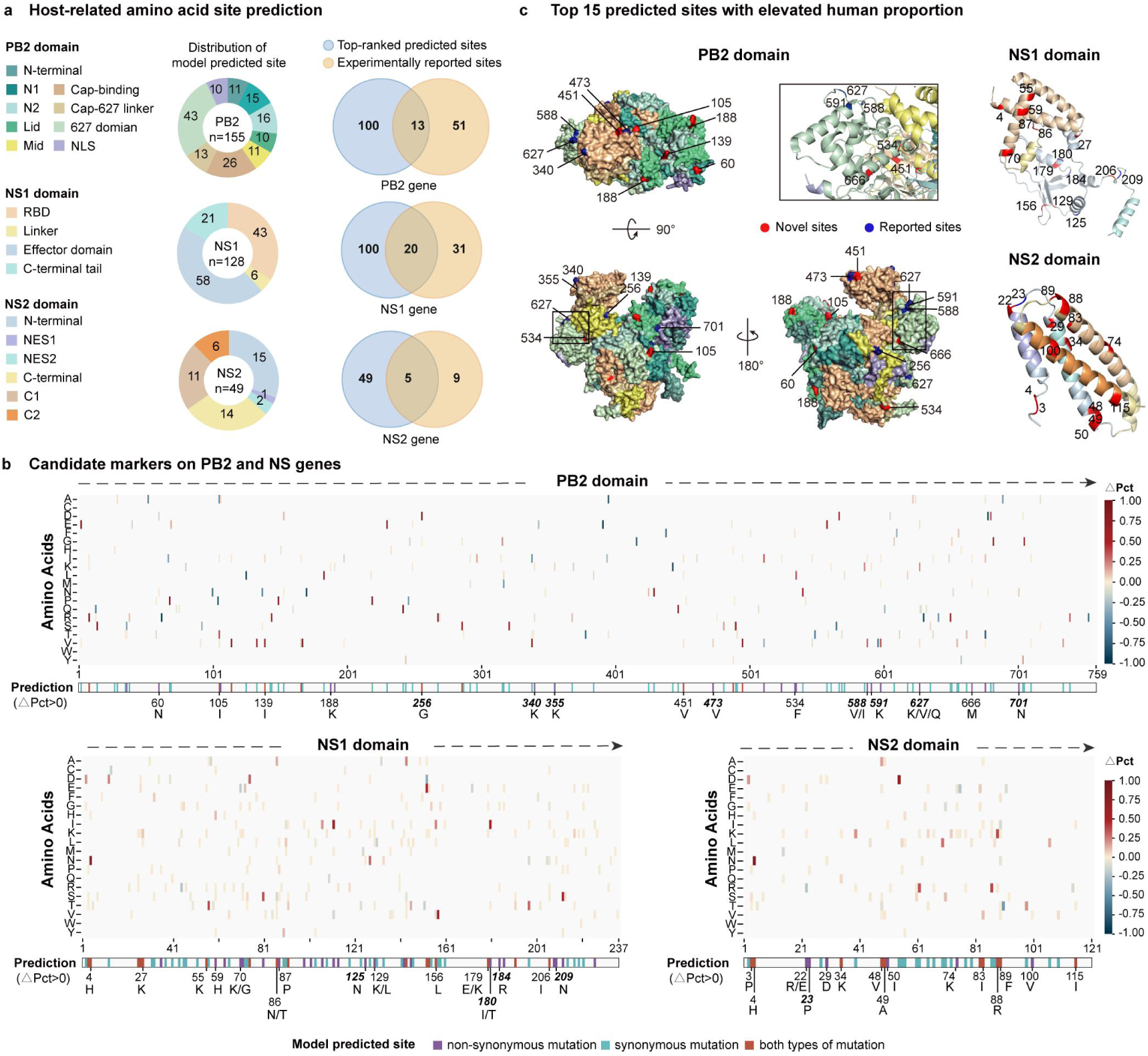
**Molecular marker identification on PB2 and NS genes for avian influenza virus spillover into humans.** a. Distribution of amino acid sites related to host tropism in PB2, NS1, and NS2 proteins identified by the model. These amino acid sites are significant for distinguishing avian and human hosts. Protein structural domains^55–57^ are marked in different colors. Venn diagrams show the overlaps between the top-rank predicted sites and the experiment reported sites. b. Candidate molecular markers on PB2, NS1, and NS2 genes. Each heatmap illustrates the difference in amino acid proportions between representatives sequences of human-infected avian influenza viruses (AIVs) and avian-derived AIVs (ΔPct) for each amino acid across all positions, with colors ranging from deep blue (ΔPct=-1) to deep red (ΔPct=1). The colored band beneath the heatmap marks different mutation types of candidate amino acid sites predicted by the model. Only sites with elevated human proportion (ΔPct>0) are shown. c. The 3D structure diagrams illustrate the top 15 amino acid sites associated with elevated human proportion (ΔPct>0) in PB2, NS1, and NS2 proteins. Previously reported sites are highlighted in blue and newly identified sites are in red. Protein domains are color-coded corresponding to those indicated in panel (a). Figures were drawn with Pymol^53^.

## Discussion

In this study, we established FluAdaX, a transformer-based deep learning framework for predicting the host adaptation of IAVs and decoding the molecular mechanisms underlying human adaptation. It provides a feasible case for applying artificial intelligence in predicting and understanding the risks of IAV cross-species transmission. FluAdaX-G makes predictions based on the integrated full-genome IAV sequences at the nucleotide level, achieving robust host classification accuracy and enabling quantitative assessment of host adaptation potential. The assessment of cross-species transmission risk in this study focused on the potential for sustained viral circulation in novel host populations, a prerequisite for IAV pandemic. Our framework achieved comparable performance to the WHO’s TIPRA framework in prioritizing zoonotic risks of major AIV strains, while surpassing traditional ML methods in host discrimination in cross-species transmission events.

Spillover score λ_sp_(γ) and adaptability score λ_ad_(X) were calculated to quantify cross-species transmission. IAV variants with elevated human adaptation potential were identified, such as H7N9, H9N2 avian IAVs, and H1N1 swine IAVs, which have been frequently studied due to thier critical role for zoonosis^8,27,28,31–34^. Swine population have been widely considered as “mixing vessels” in pandemic emergence due to their capacity to host co-infections and facilitate reassortment among influenza viruses^5,29^. Our risk assessment for SIV genome data further underscored this bidirectional transmission pattern.

Since 2021, H5 clade 2.3.4.4b viruses have experienced an unprecedented global spread via wild birds and caused spill-over infections to wide range of mammals, even humans^35^. The newly emergent H5N1 B3.13 and D1.1 genotypes, which caused outbreaks in cattle and human infections, remained adaptation to avian reservoirs according to the model prediction, which was consistent with the reinforced risk assessments by WHO classifying these viruses as primarily avian-adapted^36^. Previous studies also reported that H5N1 B3.13 viruses favored avian receptors, with slight human receptor affinity^37^. This evolutionary constraint aligns with the ecological trapping hypothesis^38^ that H5N1’s dominance in avian reservoirs, sustained through high viral loads in wild birds and environmental persistence, disincentivizes adaptation to alternative hosts. Although the emergent H5N1 viruses has not yet achieved sustainable human-to-human transmissibility, increasing mammalian adaptations (e.g., cattle-adapted PB2-631L^39^) could rapidly alter the risk profile, necessitating heightened vigilance.

We developed a two-stage screening approach by combining FluAdaX-AIV with the XGBoost algorithm, integrating the capacity of FluAdaX-AIV in host adaptation prediction and the interpretability of XGBoost to identify molecular markers associated with host tropism. PB2 and NS genes were identified as pivotal determinants of human adaptation besides surface proteins, aligned with . Most of the high-risk variants with elevated human adaptation contain H9N2 G57 internal genes (Fig. 4b; Supplementary Fig. 13), corroborating their pivotal role for novel reassortants in zoonotic emergence^8,31–34^. Recent studies focusing on the role of H9N2 in the spillover of AIVs also highlighted the significant contribution of the internal genes from H9N2 in facilitating human adaptability^40^.

Our study has several limitations. First, the model’s training on five major host taxa (avian, human, swine, canine, equine) restricted its assessment of emerging hosts. Expanding genomic data from diverse host species should be incorporated into the training set to improve generalizability. Second, while the framework performed robustly on conserved adaptive pathways (e.g., H1N1pdm09 lineage, H7N9), its sensitivity to rapidly evolving lineages remained constrained by sparse genomic surveillance data. Third, the nucleotide-level framework, though enabling broad signal detection, conflates adaptive and neutral processes. Dual-host residues (e.g., PB2-V271A^41^) critical for balancing avian fitness and human adaptation are systematically deprioritized. Future iterations must prioritize real-time surveillance integration (e.g., through GISAID updates) and functional validation via reverse genetics. Furthermore, the biological interpretation of synonymous codon biases in avian vs. human viruses remains challenging, highlights gaps in our current knowledge.

This study presents a conceptual advance in language model design for host adaptation of IAVs, offering a comprehensive framework that enables host discrimination, risk assessment of cross-species transmission, and adaptive molecular markers using exclusively IAV nucleotide sequences as input. Wang et al.^40^ investigated the early-warning signals for AIV spillover and identified associated risk factors leveraging ML methods, global epidemiological data, and phylogenetic analysis. Future efforts to enhance language models for comprehensive cross-species risk assessment of IAVs should integrate multimodal data including ecological data, spatiotemporal transmission patterns, and protein structure, etc.

FluAdaX is a dynamic, scalable and interpretable probabilistic framework that lay the foundation for full-genome modeling of IAV strains and redefine influenza pandemic preparedness through AI-driven genomics, enabling real-time monitoring of emerging IAV variants and risk assessment of their cross-species transmission potential during zoonotic outbreaks. As global zoonotic threats escalate, such AI tools will be critical for early detection of stealth adaptation in reservoir hosts, evolutionary-informed vaccine development, and targeted containment of variants exhibiting pandemic signals.

## Methods Dataset

### Endemic influenza virus dataset

A large corpus of nucleotide sequences of influenza A viruses (IAVs) were collected from EpiFlu Database on Global Initiative on Sharing All Influenza Data (GISAID), encompassing all eight segments of Influenza A viruses sampled from avian, human, swine, canine, and equine^42^ as of January 12, 2024 (a total of 1,673,576 sequences). Isolate ids, isolate names, sampling dates, and host information were extracted from the definition content of sequence file in FASTA format. We focused on high-quality coding region sequences, excluding those with incorrect labels, incomplete length, and degenerate bases. For sequences with duplicate IDs or 100% identity, a single representative was retained for each host subset. To minimize biases from different sequencing methods, we restricted our datasets to isolates collected since 2005 for model training. To balance the number of sequences across different hosts, down-sampling was applied to the human, avian, and swine subsets. The final datasets consisted of 692,386 sequences (Supplementary Fig. 1a), among which 69,062 isolates had complete genomes (Supplementary Fig. 1b).

### Zoonotic virus dataset

We constructed a second dataset of IAV with full-length coding region and complete genomes for the evaluation of host adaptability of zoonotic viruses. These sequences were obtained from GISAID Epiflu Database as of October 31, 2024 (Supplementary Fig. 1c). Sequence quality control was performed as in the first dataset, resulting in 1,674 full-genome IAV isolates included in the study (Fig 1). This dataset included full genome sequences from avian influenza virus (AIV) infections in humans (810 strains), swine influenza virus (SIV) infections in humans (130 strains), as well as those IAV strains infecting other mammals (734 strains) outside of the five host labels targeted in our model. These additional mammalian species represent a diverse range of taxa that are not typically associated with the predominant hosts of IAV but still pose a potential risk for zoonotic transmission.

### FluAdaX framework

The architecture and workflow of FluAdaX framework are show in Fig.1. The framework is initially designed to predict adaptability of the IAVs among five types of host, including human, avian, swine, canine, and quine. As illustrated in Fig.1a, the structure of the base module is a transformer-style model, with multi attention layers. We use moving average equipped gated attention (MEGA)^2^ as the backbone of FluAdaX to efficiently process extremely long sequences, in terms of computational and information extraction performance. The whole dataset of alignment-free nucleotide sequences of the IAV was partitioned into training, validation, and test sets at an 8:1:1 ratio according to the collection timeline within each host category. The outputs of FluAdaX are processed with a softmax function to generate a set of probability values (confidence level β) corresponding to the host species.

### Model architecture

We adopted the architecture of MEGA as the backbone of our model. The main idea of MEGA is using inductive biases into the attention mechanism across the tokens. The multi-dimensional damped form of exponential moving average (EMA) with learnable coefficients, and subsequently develop the moving average equipped gated attention mechanism by integrating the EMA with a variant of the single-head gated attention solved, theoretically grounded, two weaknesses simultaneously from attention mechanism: 1) weak inductive bias; and 2) quadratic computational complexity. Experimental results on various sequence modeling tasks across different data types show MEGA significantly outperforms a variety of strong baseline models, in terms of both effectiveness and efficiency.

Suppose X = {x1, x2, …, xn}∈R_n×d_ to denote a sequence of input representations with length n. Let Y = {y1, y2, …, yn}∈R_n×d_ be the sequence of output representations of each layer with the same length n as the input X. Ma et al. ^43^ assume the representations of the input and output sequences have the same dimension d. The EMA recursively calculates the output sequence Y :

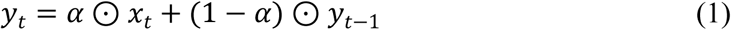

where α ∈ (0, 1)d is the EMA coefficient representing the degree of weighting decrease, and O is the element-wise product.

The gated attention mechanism in MEGA adopts the Gated Recurrent Unit (GRU^4^) and Gated Attention Unit (GAU^5^) as the backbone architectures, with an EMA-based sub-layer embedded into the calculation of the attention matrix.

### Model training and evaluation

FluAdaX-genome (FluAdaX-G) processed whole IAV genome by concatenating the alignment-free nucleotide sequences of eight segments into one single input sequence (∼13kb for each strain), with a special token(’<SEP>’) between segments. Like bert, ’<CLS>’ was attached in the first place in the sequence above which is used for representing the whole genome on classification task. We used the Unique Nucleotide (UN) Tokenizer for nucleotide sequences during model training and target prediction. The UN Tokenizer works by assigning a unique token to each unique nucleotide present in the nucleotide sequence (’A’, ’T’, ’C’, ’G’), then mapping the raw sequences into unique integer id sequences. The hyper-parameters of FluAdaX extensively referenced the raw mega setting. The hidden size is 128 and intermediate size is 256, including 2 MEGA layers. Using silu as activation function and rotary as relative positional bias. For 5 target classification, we using cross entropy loss function. The batch size of training was set to 16. The learning rate setting as 8e^−5^ and the AdamW as optimizer with one epoch warmup during optimization process. An early stop strategy was used by observing the recall rate of the validation set. FluAdaX-G was trained on A100 GPUs using a distributed computing architecture, taking about 2 days.

FluAdaX-segment (FluAdaX-S) model was meanwhile constructed for eight individual segments of IAVs using an expanded dataset. It incorporated additional IAV strains that contain partial segments, resulting in a total of 692,386 nucleotide sequences (Supplementary Fig. 1a,b), each starting with a ’<CLS>’ token. The UN Tokenizer was used as FluAdaX-G. The hidden size is 128 and intermediate size is 256, including 4 MEGA layers. Other settings were the same as FluAdaX-G. FluAdaX-S was trained on A100 GPUs using a distributed computing architecture, taking about 20 hours.

Model performance was assessed using metrics as follows. The accuracy and recall was used for the classification task. For observation host adaptability or infection portability performance, AUC result was also calculated.

### Risk assessment based on FluAdaX-G Calculate baseline index for IAV host adaptability based on endemic strain predictions

Based on the FluAdaX-G’s full-genome prediction of host adaptability, we characterized the host adaptation level within 6,904 IAV strains circulating among human, avian, swine, canine, and equine (up to January 2024). This approach considered the host adaptation mechanism from a genome-wide perspective, and established a baseline for the adaptability levels of circulating strains to different hosts. Based on this baseline, we can further identify the relative adaptability of newly emerging IAV strains to alternative hosts compared to their sampling origin by calculating spillover score λ_sp_(γ) and adaptability score λ_ad_(X).

Th spillover score λ_sp_(γ) quantifies the overall spillover risk of IAV strain from the sampling host, defined as:

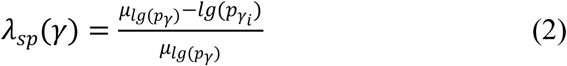

where *p_γi_* represents the predicted probability of strain *i* infecting host γ (human, swine, avian, canine, equine), and μlg(p_γ_) is the mean log-transformed probability of infection for host γ across all samples within specific host. To ensure comparability of spillover score across five host species, λ_sp_(γ) was normalized as λ’_sp_(γ), fomulated as:

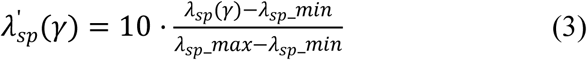

where λ_sp_(γ)) is the original spillover score for each sample. λ_sp__min and λ_sp__max are the minimum and maximum λ_sp_(γ) across the five hosts. The final result for endemic IAVs falls within the range [0, 10].

The adaptability score λ_ad_(X) quantifies the adaptability of IAV strains circulating in a certain host γ to specific host X, the risk function can be formulated as:

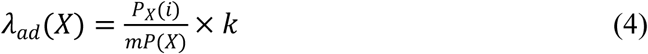

where *P_X_(i)* is the predicted probability of strain *i* for host *X* (human, swine, avian, canine, equine). *mP(*X) is the median infection probability for host *X* within the IAV strains sampling from the same host species, which represents the average adaptability of circulating IAV strains within each host. *k* is an adjustment parameter that controls the sensitivity of the adaptability score. In the calculation of adaptability to the host of origin, *k*=1; when considering adaptability to other hosts, *k*=0.001. For each virus strain, an adaptability score is calculated for each of the five host types, denoted as λ_ad_(human), λ_ad_(swine), λ_ad_(avian), λ_ad_(canine), λ_ad_(equine). To ensure comparability of adaptability scores among IAV strains across different host species, we normalized the adaptability scores for each of the five host types (human, swine, avian, canine, equine). The normalized adaptability score λ’_ad_(X) was calculated using the following equation:

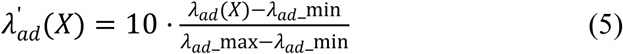

where λ_ad__min is the minimum value of λ_ad_(X) across all host types, λ_ad__max is the maximum value of λ_ad_(γ) across all host types. The normalized adaptability scores are within the range of 0 to 10.

Finally, we use the normalized spillover scores λ’_sp_(γ), and adaptability scores λ’_ad_(X) to assess the relative adaptability of IAV strains to different hosts. For simplification, these normalized scores are uniformly denoted as λ_sp_(γ) and λ_ad_(X) in the article and below.

### Assess the human adaptability of zoonotic viruses

We evaluate the adaptability of cross-species cases to humans using the zoonotic virus dataset. The adaptability scores for human-infected AIVs and other mammals-infected AIVs are calculated relative to avian endemic strains, while the adaptability scores for human-infected with SIVs are calculated relative to swine endemic strains. This approach quantifies the risk of human adaptation after cross-species transmission of IAV strains.

### Comparative analysis

To benchmark the performance of FluAdaX series models for IAV host adaptability characterization, we compared it with traditional machine learning (ML) methods. These included Support Vector Machine (SVM)^44^, Gradient Boosting Machine (GBM)^45^, Decision Tree-based Adaptive Boosting (AdaBoost)^46^, and multi-layer perceptron (MLP)^47^. All ML analyses were performed with Python using the Scikit-learn package (v1.3.0) under uniform training protocols. We used the same datasets to train our model and these alternative methods to ensure the comparability of our results. All models processed concatenated full-genome sequences and individual gene segments, with identical splits of training (80%), validation (10%), and test (10%) sets chronologically partitioned. Nucleotide sequences were preprocessed into 3-mer frequency vectors for these four algorithms. For SVM, a linear kernel with C=1 and one-vs-rest strategy leveraged LIBLINEAR optimization to handle high-dimensional genomic data, prioritizing margin maximization in the 3-mer feature space. The GBM ensemble combined 300 gradient-boosted decision trees (max depth=4, learning rate=0.01), iteratively minimizing multinomial deviance through positional 3-mer counts that encoded reading frame-specific biases. AdaBoost employed decision tree stumps (max depth=4) as weak learners, sequentially reweighting misclassified sequences over 300 iterations using the SAMME algorithm. The MLP architecture, optimized via randomized hyperparameter search, adopted a single hidden layer (375 ReLU units, L2 regularization α=0.0034) trained with Adam optimizer (learning rate=0.01). Performance of these ML algorithms was evaluated using Area Under the ROC Curve (AUC), recall, and accuracy metrics on held-out test sets spanning 6,904 endemic IAV strains and an additional sets of 1,674 zoonotic IAV strains (Fig.2a; Supplementary Fig. 1b,c).

For the assessment of human adaptability of zoonotic IAVs, we further compared FluAdaX-G results against TIPRA evaluation^11^. Major clade or lineage of viruses that have reported infections in humans, including H7N9, clade 2.3.4.4 H5Nx, clade 2.3.2.1c H5N1, clade 2.3.4.4 H5N6, H9N2 Y280 lineage, and H9N2 G1 lineage were evaluated. Mean point estimate scores for human infection assessed by TIPRA were compared with the mean values and upper-bound estimates of human adaptability scores assessed based on the output of FluAdaX-G. Pearson correlation analysis was used to assess the consistency between the results of the two methods.

### Molecular marker analysis

#### Identify adaptation signals based on model prediction

We utilized multiple sequence alignments (MSAs) of 113,722 avian-derived and 5,585 human-infected avian influenza viruses (AIVs) to identify the key molecular determinants associated with host tropism between avian and human (Supplementary table 6). Sequences were aligned using MAFFT version 7.222^48^ and manually adjusted to correct frameshift errors in MEGA v7.0^49^. Segment-specific models were constructed using the same architecture as FluAdaX for host discrimination with each MSA of the eight viral segments. For the HA and NA segments, H9 and N2 genes were used in this study. The avian-derived sequences of HA and NA were downsampled using CD-HIT^50^ to maintain a ratio of approximately 10:1 relative to human sequences. The MSAs of each segment from avian-derived and human-infected AIVs were divided chronologically at a ratio of 8:1:1 for training, validation, and test sets. The output of models were a set of probability values for avian and human infection, indicating the likelihood of a given sequence being adapt to avian/human. Next, a two-step approach were applied to identify critical nucleotide positions for host discrimination in nucleotide sequences.

Initially, we used the influence function to select the top 600 representative gene segments that had the greatest impact on the model’s predictive performance (Supplementary Fig. 14). The influence function, a well-established technique in robust statistics, quantifies the effect of small perturbations in the training data on the model’s output^51^. Specifically, it measures the change in the model’s prediction caused by perturbing individual data points, enabling the identification of influential observations that significantly contribute to the overall model behavior. By applying the influence function, we were able to isolate the gene segments that were most informative for the task at hand, ensuring that only the most relevant data were used for subsequent analysis.

Following this, we employed the XGBoost algorithm^52^, a gradient boosting framework renowned for its effectiveness in classification tasks, to pinpoint the key nucleotide positions within the selected sequences. To prepare the data for model training, each nucleotide position was encoded using a set of five features representing the presence of adenine (A), thymine (T), cytosine (C), guanine (G), and gaps (”-”). This encoding enabled the XGBoost model to evaluate the importance of each nucleotide position in relation to host discrimination. By training the model, we obtained feature importance scores, which allowed us to identify the nucleotide positions and their corresponding bases that were most critical for distinguishing between different hosts. Nucleotides with non-zero feature important score were selected as candidate markers for further analysis. The combination of the influence function and XGBoost provided a powerful framework for identifying biologically significant sequence motifs and positions, offering valuable insights for further sequence-based analysis and model refinement.

#### Analyze key mutations related to host tropism

Codon compositions significantly associated with host tropism were analyzed and these nucleotide positions were then transformed into amino acid position, which can be precisely interpreted corresponding to other experimental studies. Published data were analyzed to screen for mutations experimentally validated for altered mammalian adaptation. Codon distribution at each site were analyzed for different hosts within the top 600 representative sequences of the each of dominant internal genes (Supplementary Fig. 14). We calculated the distribution proportion of codons at each site in avian and human hosts, denoted as *Ci*(Avian)% and *Ci*(Human)%, respectively. Subsequently, we calculated the difference of codon compositions between human and avian at each site, formulated as ΔPct=*Ci*(Human)% – *Ci*(Avian)%. The ΔPct values range from -1 to 1, with ΔPct>0 indicating a preference for human-biased codon at that site while ΔPct<0 indicating the opposite. We then filtered positions and codon compositions with ΔPct>0 in our model prediction. Finally, we selected sites with non-synonymous mutations of AIVs with human-biased codon composition and their corresponding amino acids as priorities. These amino acid sites were ranked according to the feature importance scores assessed by the model. All of the predicted amino acid positions were mapped to the protein domain, and the top 15 sites with non-synonymous mutations (meanwhile ΔPct>0) were visualized on the 3D protein structure through PyMoL v3.0.4^53^.

### Phylogenetic analyses

The adaptability scores were correlated with the phylogeny of IAV to explore the underlying mechanism of human adaptation. We reconstructed maximum-likelihood (ML) phylogenies of all segments from H1N1 IAV test set by using FastTree version 2.1.11^54^. The datasets were downsampled based on the HA sequence similarity in each host using CD-HIT^50^. For human H1N1 strains with 99% similarity in HA sequences, one representative was retained, so do the swine strains. Two avian H1N1 AIVs and human-infected H1N1 SIVs after 2019 were retained, resulting in a total of 147 full genome strains. All gene segments of these viruses, together with those of four reference H1N1 strains, were employed for the tree reconstruction. The resulting ML tree was classified into divergent lineages. Spillover scores λ_sp_(γ), and adaptability scores λ_ad_(X) were labeled aside the tree. ML trees were also reconstructed for gene sequences of human-infected AIVs and genotype analysis was then conducted for these strains.

## Data availability

The IAV nucleotide sequence data used for model generation and evaluation are available from the GISAID EpiFlu Database (https://gisaid.org/).

## Code availability

The deep-learning models were developed and deployed using standard model libraries and the PyTorch framework. Code is available at GitHub (https://github.com/MiaJY-Yang/FluAdaX/).

## Acknowledgements

We gratefully acknowledge the authors and laboratories for sharing the influenza virus sequences in GISAID database. This study was supported by the National Key Research and Development Program of China (2021YFC2300100), Non-profit Central Research Institute Fund of Chinese Academy of Medical Sciences (2022-RC310-02), and the National Nature Science Foundation of China (81961128002_82341118).

## Author contributions

J.Y., Z.L., and Y.S. conceptualized and designed the study. P.F. and J.Y. prepared the data and designed the FluAdaX framework. P.F. and J.Y. evaluated the performance of FluAdaX series models and other machine learning methods. J.Y. performed interpretation of results and molecular analyses. J.Y., and P.F. wrote the original draft of the manuscript. J.Y., L.Y., J.L.,Z.L. and W.Z. made critical revision of the manuscript. All authors discussed the results and reviewed the paper.

