## Supplementary material for "Using Artificial Intelligence to Assess Cross-Species Transmission Potential of Influenza A Virus": Figure legends of Supplementary Figures

**Figure legends**

**Supplementary Fig.1 Dataset used in AI model. a.** Number of nucleotide sequences of influenza A viruses from epidemic hosts used in this study. Original nucleotide sequences of influenza A viruses were downloaded on GISAID EpiFlu Database up to January 12, 2024. **b.** Data division for IAV genomes used in full-genome (FluAdaX-G) and segment model (FluAdaX-S). **c.** Zoonotic IAV datasets tested in this study. Full genome of IAV was concatenated as a long sequence for each strain.

**Supplementary Fig.2 Risk assessment of endemic influenza A viruses (IAVs) from various hosts evaluated by FluAdaX-G. a.** Confidence levels of FluAdaX-G predictions for the hosts of endemic IAV strains. From left to right, the boxplot and scatter plot jointly illustrate the distribution of probabilities of the strains predicted to be human, swine, avian, canine, and equine, respectively, scaled from 0 to 1. The colors of the scatter points represent the origin of the strains, including human, swine, avian, canine, and equine. The upper and lower whiskers of the boxplot represent the upper and lower quartiles, respectively, with the line inside indicating the median. **b.** Adaptability scores evaluated for endemic IAV strains. From left to right, the boxplot and scatter plot jointly illustrate the distribution of adaptability risk to human, swine, avian, canine, and equine, scaled from 0 to 10. The colors of the scatter points represent the origin of the strains, including human, swine, avian, canine, and equine. The upper and lower whiskers of the boxplot represent the upper and lower quartiles, respectively, with the line inside indicating the median.

**Supplementary Fig.3 Spillover score of whole-genome dataset and each segment of endemic influenza A viruses (IAVs).** The boxplot and scatter plot jointly illustrate the distribution of fitness loss to avian, canine, equine, swine, and human, scaled from 0 to 10. The colors of the scatter points represent the origin of the strains. The upper and lower whiskers of the boxplot represent the upper and lower quartiles, respectively, with the line inside indicating the median.

**Supplementary Fig.4 Phylogenetic tree of HA genes of H1N1 influenza A viruses tested in FluAdaX-G (n=151).** Human H1N1 strain are in blue, while reported human-infected SIVs are in violet. Swine H1N1 strain and avian H1N1 strain are denoted in orange and red, respectively. Reference sequences are marked with dot. Branch lengths are scaled according to the numbers of substitutions per site. Branch support values of selected nodes are shown. Lineages are labelled with vertical lines on the right.

**Supplementary Fig.5 Phylogenetic tree of NA genes of H1N1 influenza A viruses tested in FluAdaX-G (n=151).** Human H1N1 strain are in blue, while reported human-infected SIVs are in violet. Swine H1N1 strain and avian H1N1 strain are denoted in orange and red, respectively. Reference sequences are marked with dot. Branch lengths are scaled according to the numbers of substitutions per site. Branch support values of selected nodes are shown. Lineages are labelled with vertical lines on the right.

**Supplementary Fig.6 Phylogenetic tree of PB2 genes of H1N1 influenza A viruses tested in FluAdaX-G (n=151).** Human H1N1 strain are in blue, while reported human-infected SIVs are in violet. Swine H1N1 strain and avian H1N1 strain are denoted in orange and red, respectively. Reference sequences are marked with dot. Branch lengths are scaled according to the numbers of substitutions per site. Branch support values of selected nodes are shown. Lineages are labelled with vertical lines on the right.

**Supplementary Fig.7 Phylogenetic tree of PB1 genes of H1N1 influenza A viruses tested in FluAdaX-G (n=151).** Human H1N1 strain are in blue, while reported human-infected SIVs are in violet. Swine H1N1 strain and avian H1N1 strain are denoted in orange and red, respectively. Reference sequences are marked with dot. Branch lengths are scaled according to the numbers of substitutions per site. Branch support values of selected nodes are shown. Lineages are labelled with vertical lines on the right.

**Supplementary Fig.8 Phylogenetic tree of PA genes of H1N1 influenza A viruses tested in FluAdaX-G (n=151).** Human H1N1 strain are in blue, while reported human-infected SIVs are in violet. Swine H1N1 strain and avian H1N1 strain are denoted in orange and red, respectively. Reference sequences are marked with dot. Branch lengths are scaled according to the numbers of substitutions per site. Branch support values of selected nodes are shown. Lineages are labelled with vertical lines on the right.

**Supplementary Fig.9 Phylogenetic tree of NP genes of H1N1 influenza A viruses tested in FluAdaX-G (n=151).** Human H1N1 strain are in blue, while reported human-infected SIVs are in violet. Swine H1N1 strain and avian H1N1 strain are denoted in orange and red, respectively. Reference sequences are marked with dot. Branch lengths are scaled according to the numbers of substitutions per site. Branch support values of selected nodes are shown. Lineages are labelled with vertical lines on the right.

**Supplementary Fig.10 Phylogenetic tree of MP genes of H1N1 influenza A viruses tested in FluAdaX-G (n=151).** Human H1N1 strain are in blue, while reported human-infected SIVs are in violet. Swine H1N1 strain and avian H1N1 strain are denoted in orange and red, respectively. Reference sequences are marked with dot. Branch lengths are scaled according to the numbers of substitutions per site. Branch support values of selected nodes are shown. Lineages are labelled with vertical lines on the right.

**Supplementary Fig.11 Phylogenetic tree of NS genes of H1N1 influenza A viruses tested in FluAdaX-G (n=151).** Human H1N1 strain are in blue, while reported human-infected SIVs are in violet. Swine H1N1 strain and avian H1N1 strain are denoted in orange and red, respectively. Reference sequences are marked with dot. Branch lengths are scaled according to the numbers of substitutions per site. Branch support values of selected nodes are shown. Lineages are labelled with vertical lines on the right.

**Supplementary Fig.12 Adaptability score of zoonotic influenza A viruses (IAVs). a.** Comparison of adaptation risk of avian influenza viruses (AIVs) to human across avian and human infection cases. **b.** Comparison of adaptation risk of swine influenza viruses (SIVs) to human across swine and human infection cases. **c.** Adaptability score of zoonotic IAVs. From left to right, the boxplot and scatter plot jointly illustrate the distribution of adaptability risk to human, swine, avian, canine, and equine for different IAV origins. The colors of the scatter points represent the origin of the strains, including human-infected AIV, human-infected SIV, cattle-infected AIV, and other mammal-infected IAV. **d.** Comparative adaptation risk of zoonotic IAV subtypes to humans. The boxplot and scatter plot jointly illustrate the distribution of adaptation risks across different IAV subtypes. The colors of the scatter points represent the origin of the strains, including human-infected SIV, human-infected AIV, and other mammal-infected IAV. The upper and lower whiskers of the boxplot represent the upper and lower quartiles, respectively, with the line inside indicating the median. Differences between subtypes were compared using Mann-Whitney U tests, with * indicating *P*<0.05, ** indicating *P*<0.01, and *** indicating *P*<0.001. The black triangle above specific box plot denotes statistically significant differences between this subtype and all others.

**Supplementary Fig.13 Human adaptation risk assessment of human-infected AIV strains.** Strain names of representatives were shown alongside their internal gene cassette labeled with colored dots, with bold italic strain names indicating those of particular concern or recently prevalent. The boxplot and scatter plots on the right represent the human adaptability score λ_ad_(X) corresponding to those strains. The upper and lower whiskers of the boxplot represent the upper and lower quartiles, respectively, with the line inside indicating the median.

**Supplementary Fig.14 Prediction of molecular markers associated with host tropism between avian and human.** a.Top 600 representative gene segments for model predictive performance. Representatives of PB2 and NS genes were selected using the influence function, respectively. The number of diverse subtypes among human-infected AIVs and avian-derived AIVs were shown, respectively. b. The codon positions of predicted nucleotides related to host tropism between avian and human. c. The bar graph illustrates the overall distribution of synonymous and non-synonymous mutations associated with host tropism predicted by the model across PB2, NS1, and NS2 genes at the codon level. d-f. present the detailed information of synonymous and non-synonymous mutations in each protein domain of the PB2 (d), NS1 (e), and NS2 (f) genes, respectively. These molecular markers were identified on top representatives gene segments for prediction performance using the XGBoost models.

.
