## Supplementary Fig. 1 for "Using Artificial Intelligence to Assess Cross-Species Transmission Potential of Influenza A Virus"

a

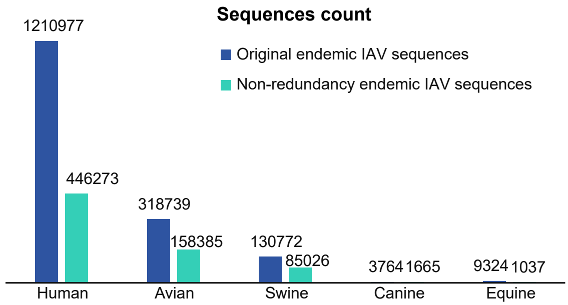


b

**Endemic infliuenza A virus genomes used in this study**

| Segment | Dataset | Human | Swine | Avian | Canine | Equine | Total |
|---|---|---|---|---|---|---|---|
| Whole genome | train | 35701 | 5513 | 13771 | 194 | 74 | 55253 |
| | valid | 4462 | 689 | 1721 | 24 | 9 | 6905 |
| | test | 4463 | 690 | 1722 | 19 | 10 | 6904 |
| HA | train | 78138 | 11246 | 23914 | 210 | 167 | 113675 |
| | valid | 8184 | 1618 | 2466 | 27 | 22 | 12317 |
| | test | 8376 | 1482 | 2448 | 27 | 50 | 12383 |
| NA | train | 61270 | 10578 | 17564 | 186 | 162 | 89760 |
| | valid | 6034 | 1497 | 2250 | 20 | 18 | 9819 |
| | test | 6933 | 1493 | 2071 | 16 | 40 | 10553 |
| PB2 | train | 47691 | 6582 | 17357 | 162 | 86 | 71878 |
| | valid | 6891 | 779 | 2386 | 26 | 13 | 10095 |
| | test | 6334 | 795 | 2163 | 18 | 3 | 9313 |
| PB1 | train | 44278 | 6456 | 15015 | 157 | 84 | 65990 |
| | valid | 6105 | 784 | 2363 | 26 | 12 | 9290 |
| | test | 5090 | 786 | 2198 | 18 | 3 | 8095 |
| PA | train | 46080 | 6710 | 16995 | 165 | 90 | 70040 |
| | valid | 7074 | 800 | 2407 | 17 | 15 | 10313 |
| | test | 6846 | 800 | 2673 | 17 | 2 | 10338 |
| NP | train | 35178 | 6314 | 15610 | 154 | 80 | 57336 |
| | valid | 4645 | 766 | 2093 | 18 | 9 | 7531 |
| | test | 4422 | 786 | 2068 | 21 | 3 | 7300 |
| MP | train | 27369 | 7099 | 13439 | 113 | 85 | 48105 |
| | valid | 2892 | 647 | 1204 | 17 | 9 | 4769 |
| | test | 3362 | 716 | 1365 | 11 | 2 | 5456 |
| NS | train | 17016 | 5699 | 14150 | 206 | 75 | 37146 |
| | valid | 2801 | 694 | 1586 | 18 | 5 | 5104 |
| | test | 3264 | 728 | 1771 | 15 | 2 | 5780 |

c

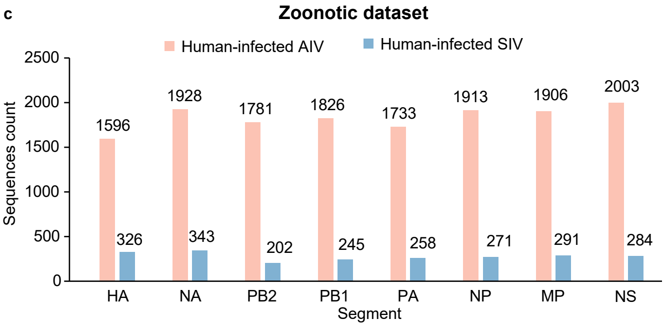
