## Supplementary figures and images for "Using Artificial Intelligence to Assess Cross-Species Transmission Potential of Influenza A Virus"

### Supplementary Fig. 2

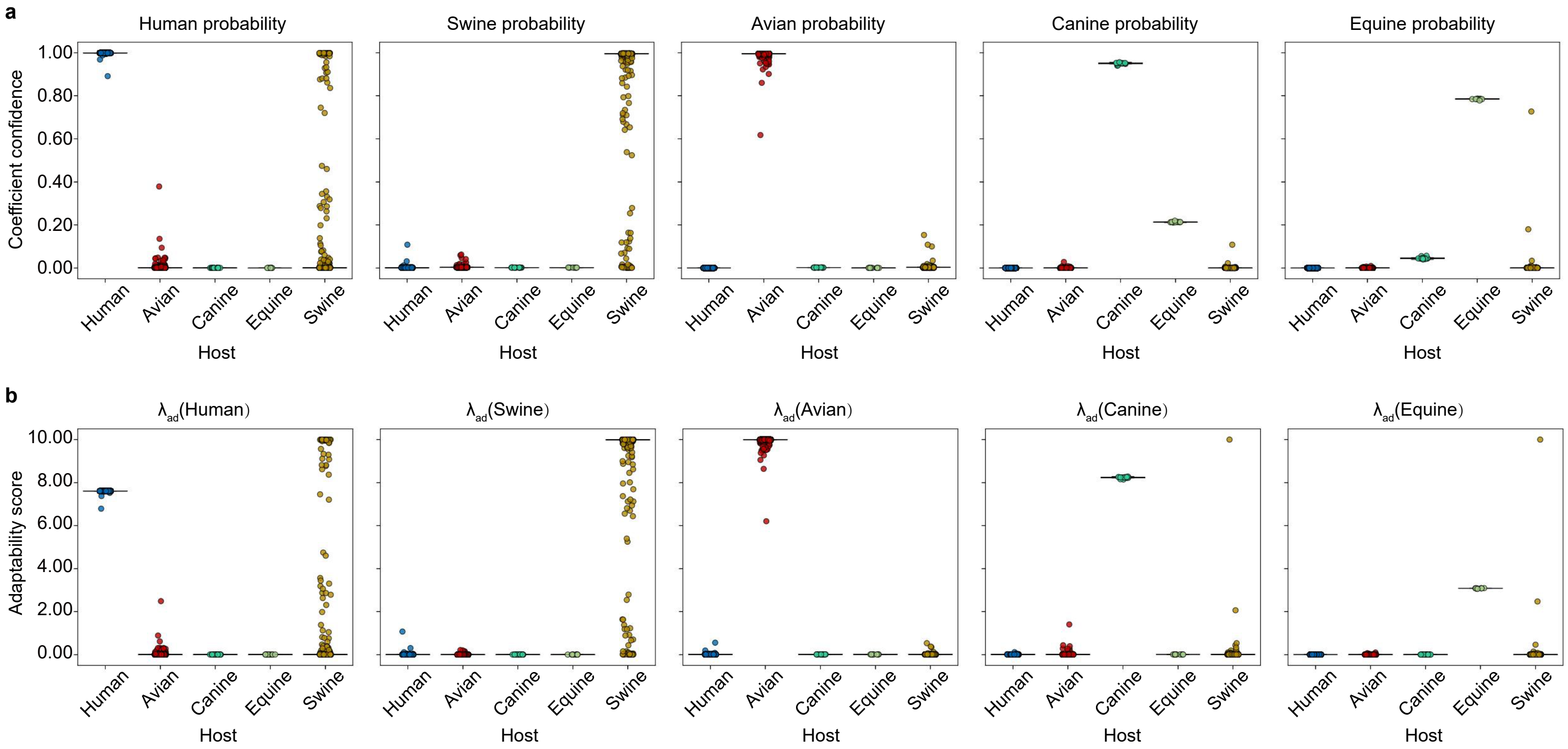

### Supplementary Fig. 3

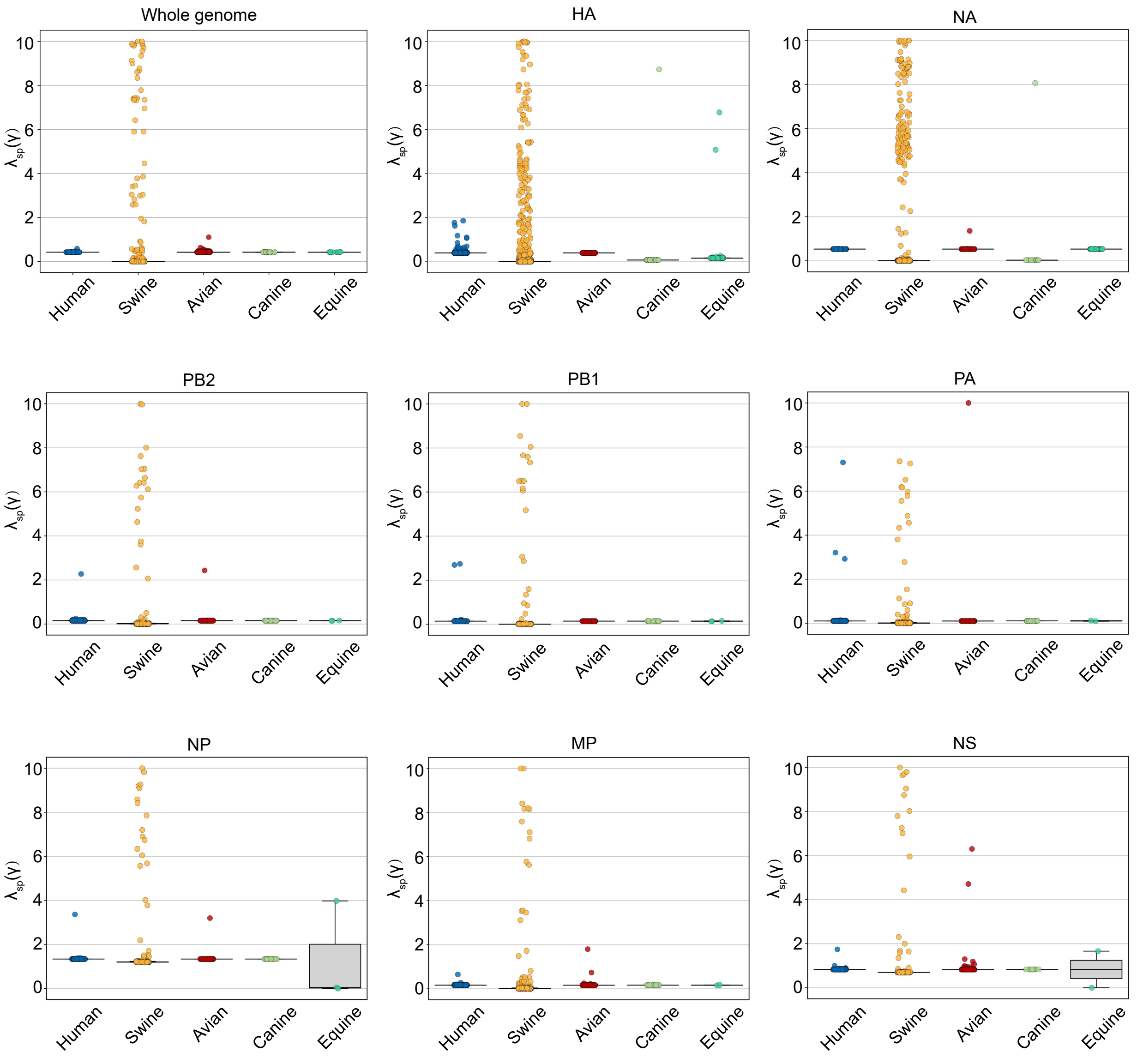

### Supplementary Fig. 4

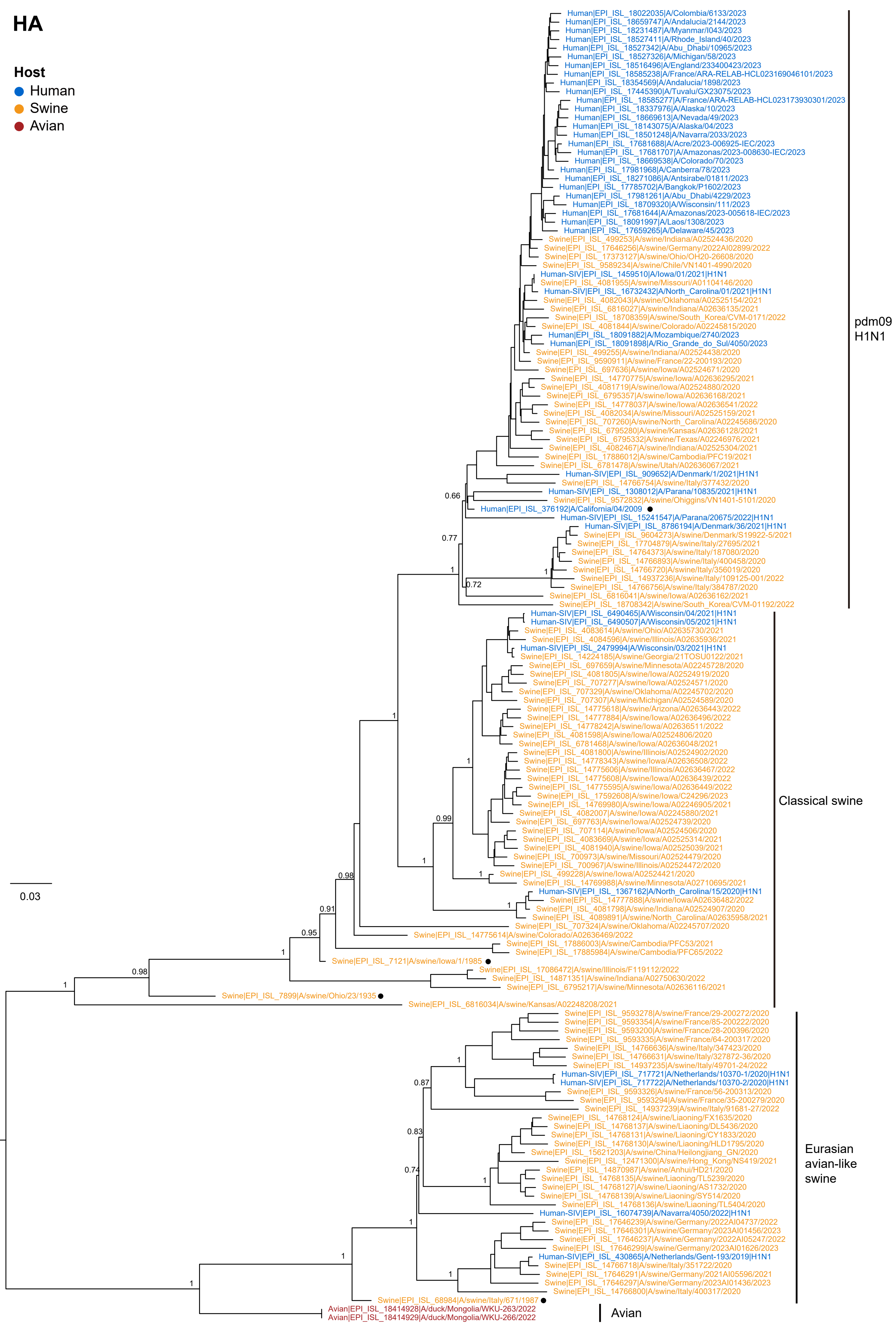

### Supplementary Fig. 5

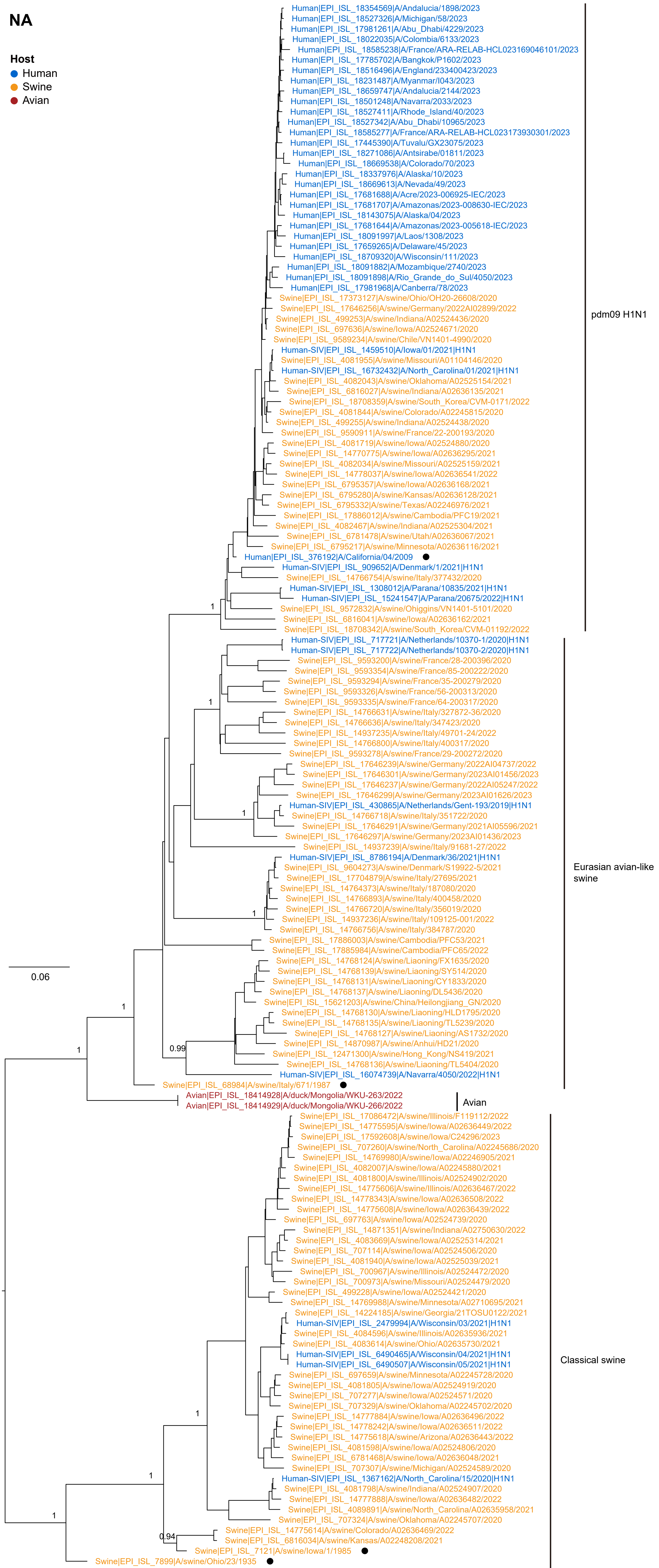

### Supplementary Fig. 6

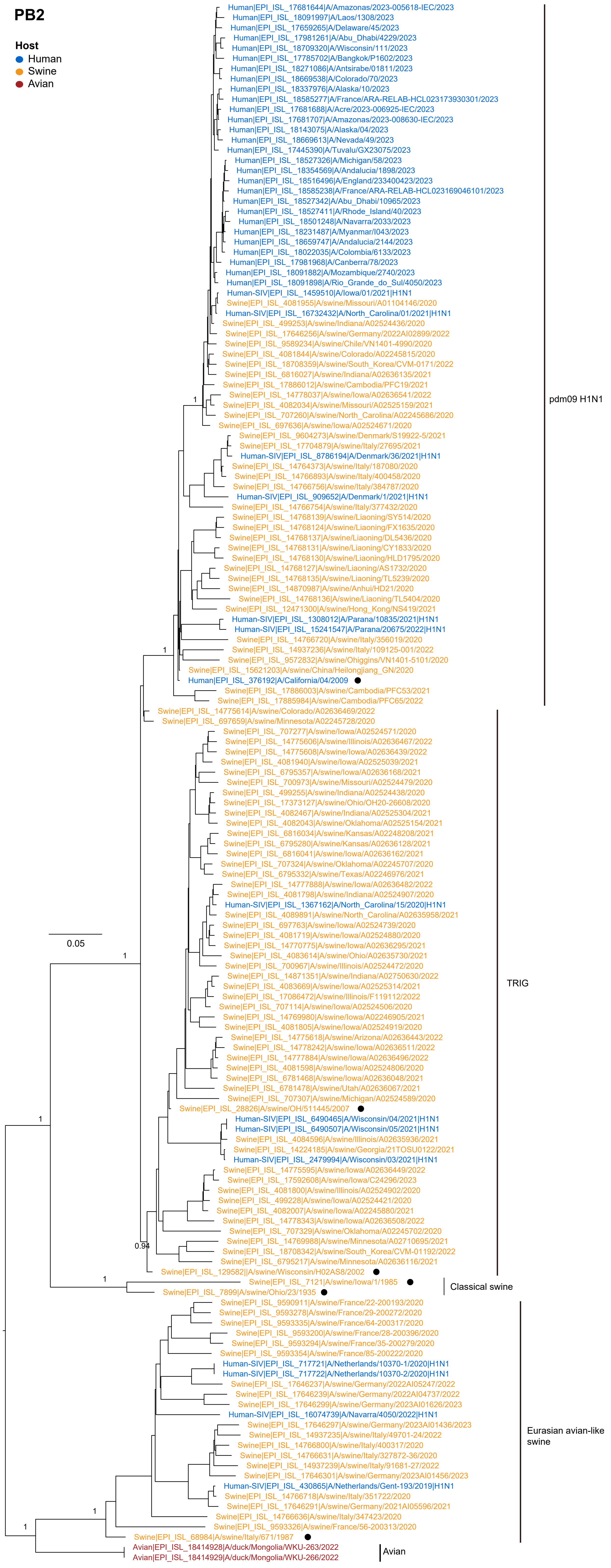

### Supplementary Fig. 7

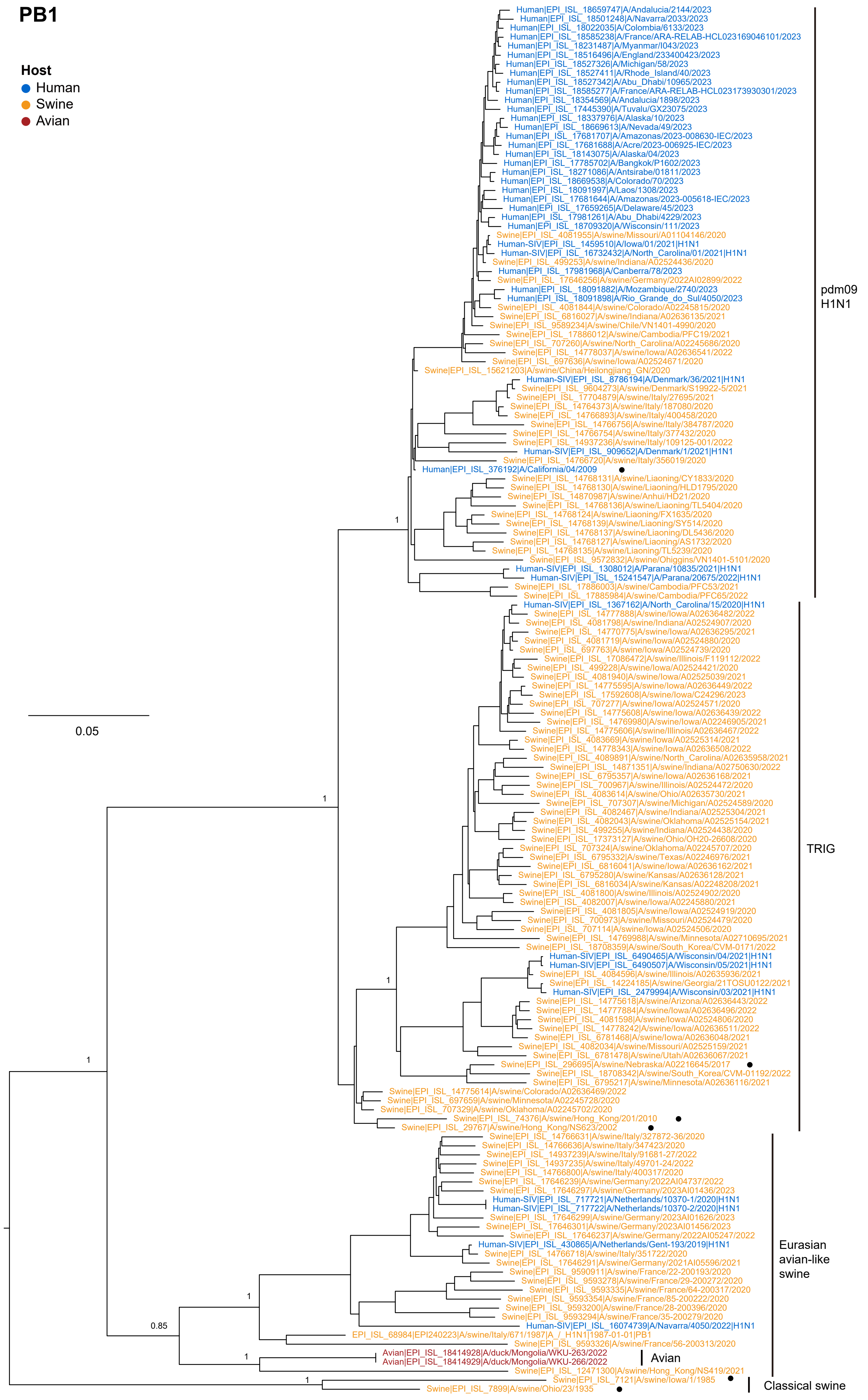

### Supplementary Fig. 8

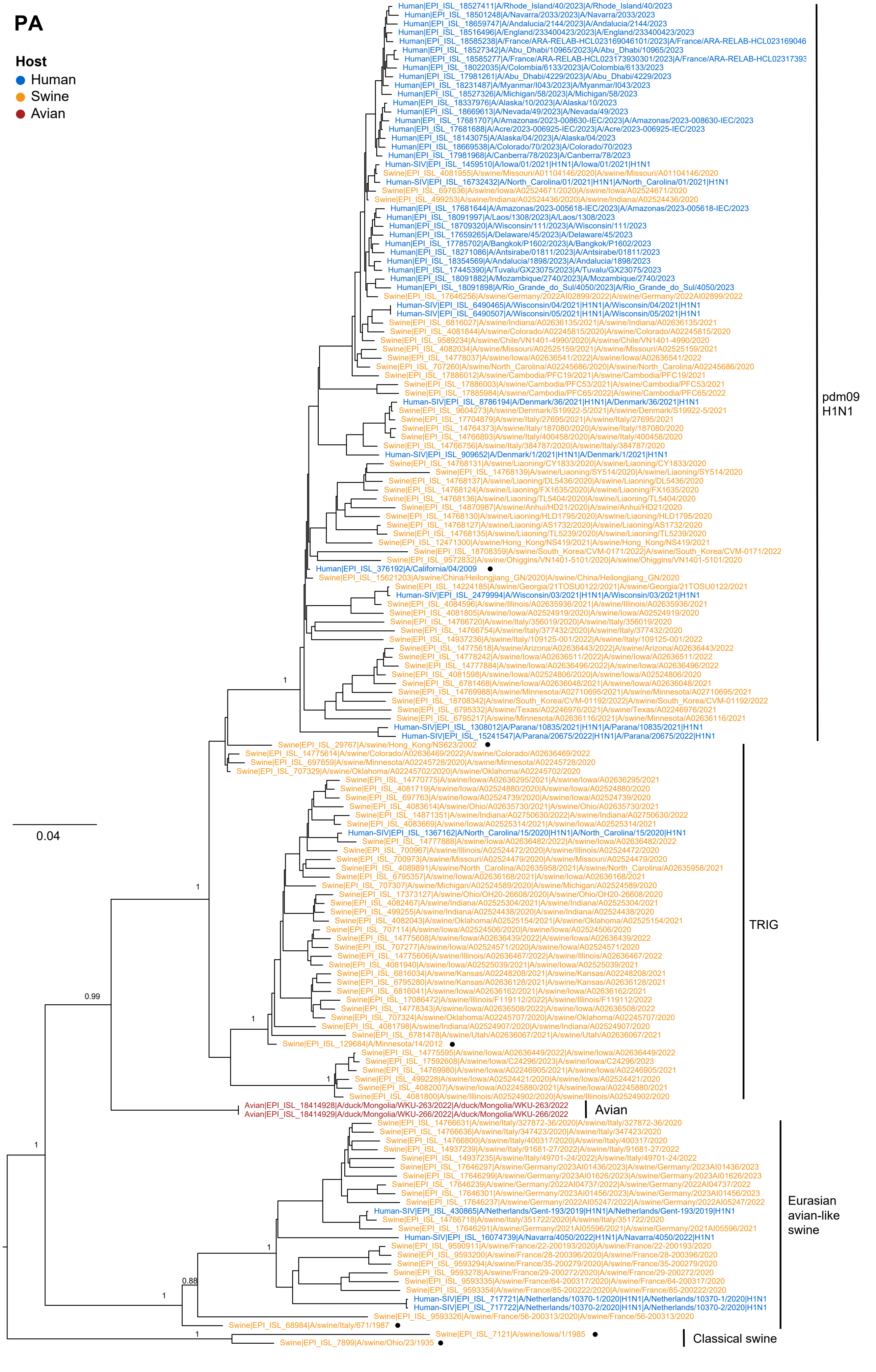

### Supplementary Fig. 9

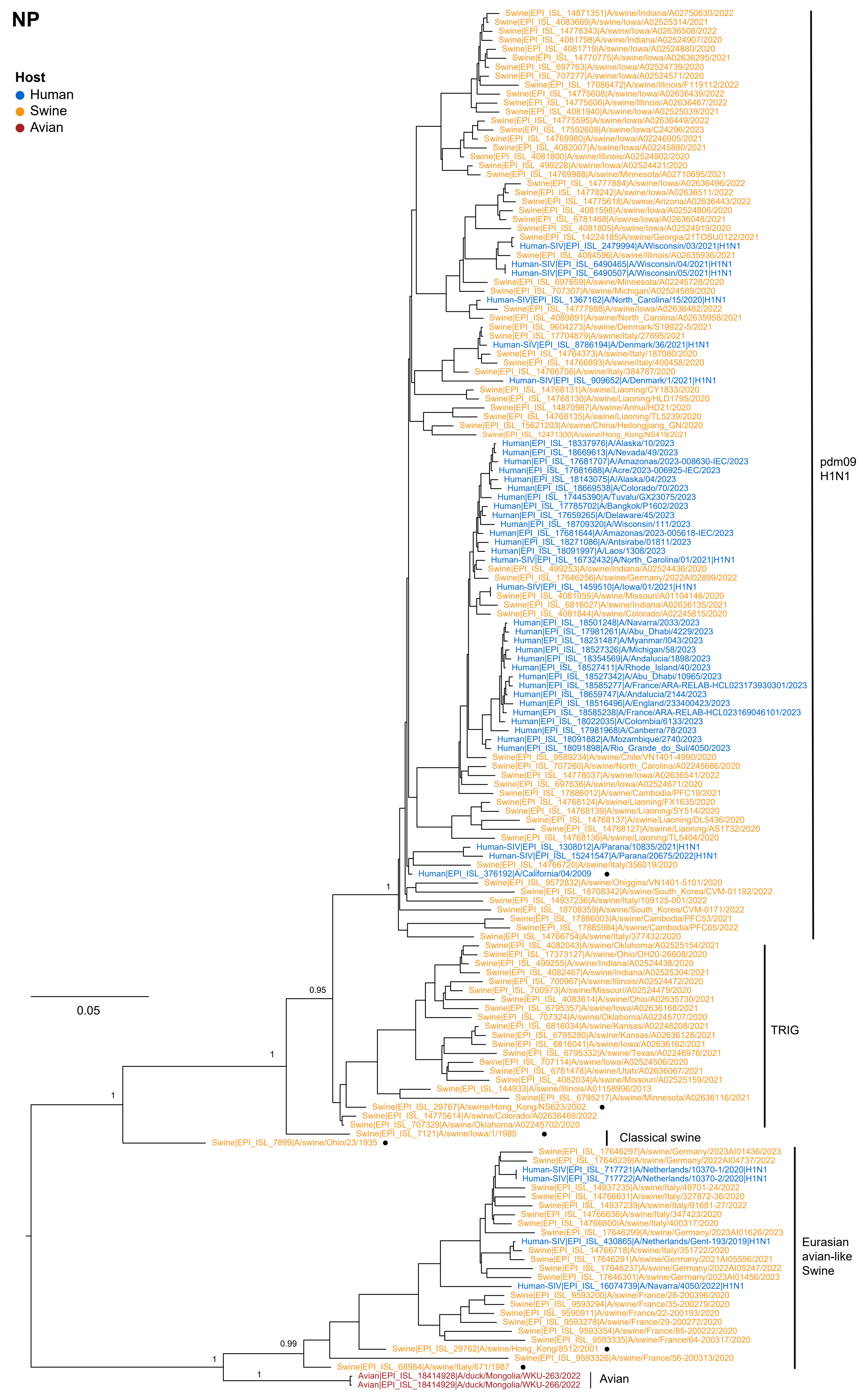

### Supplementary Fig. 10

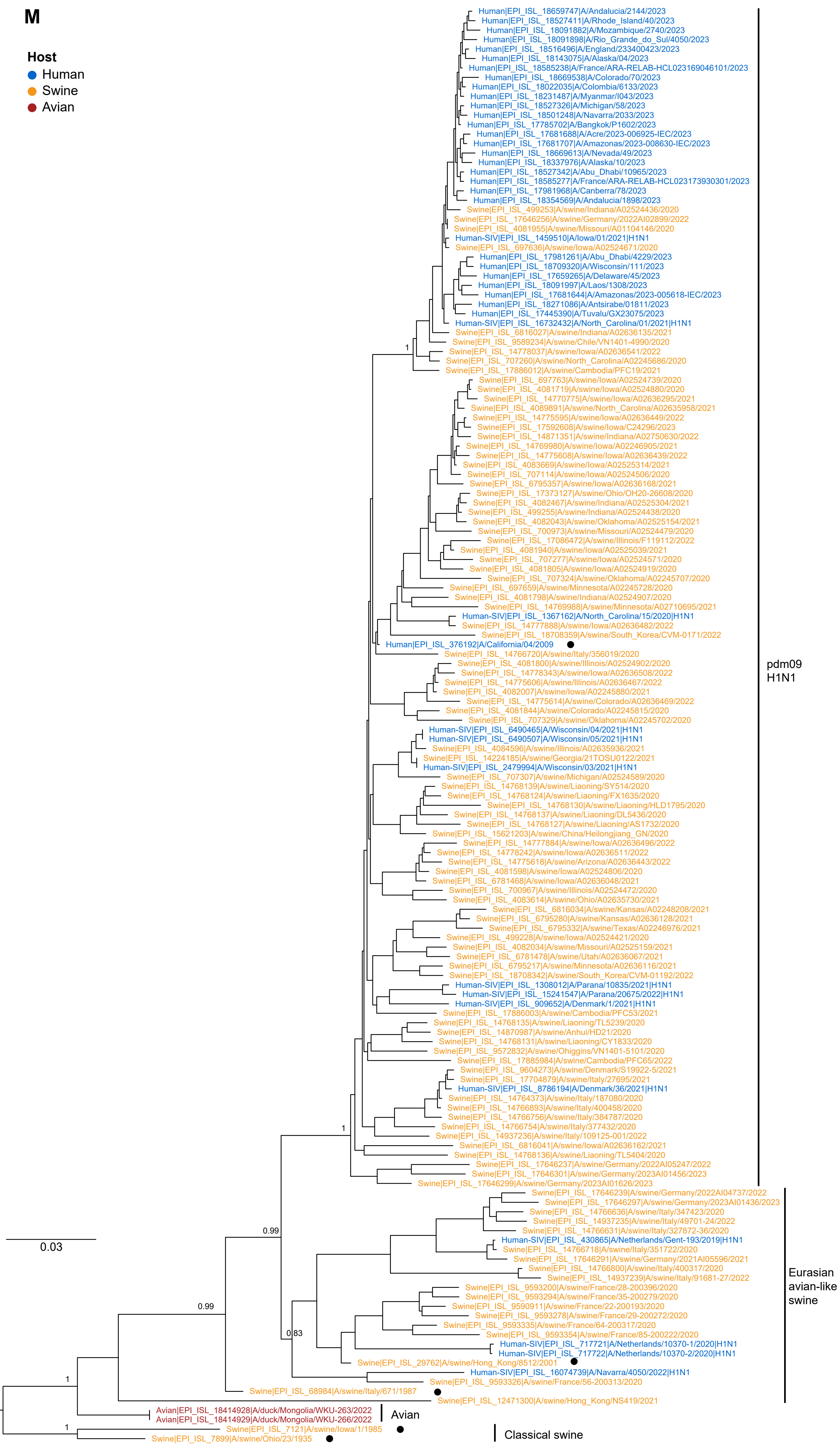

### Supplementary Fig. 11

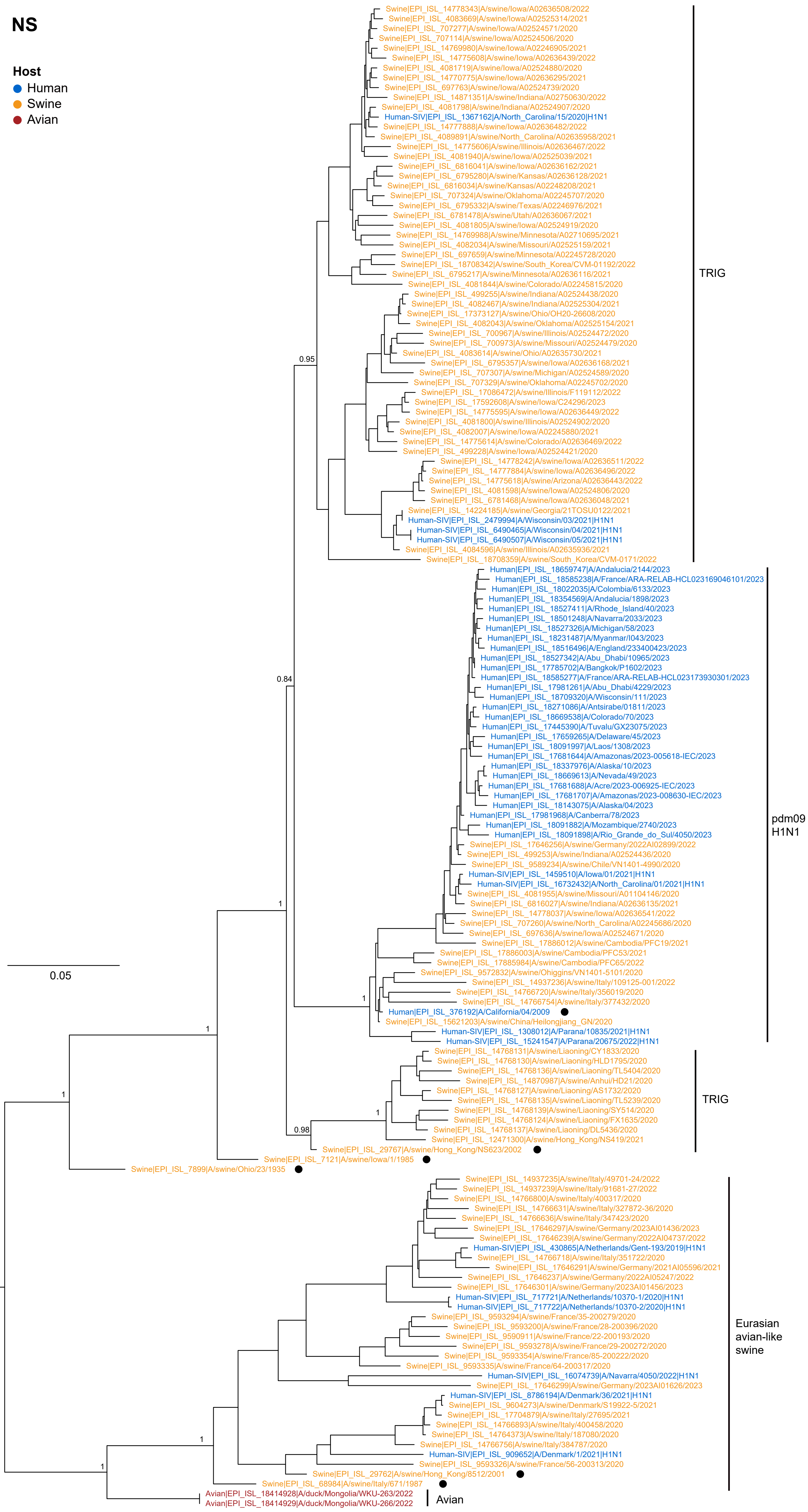

### Supplementary Fig. 12

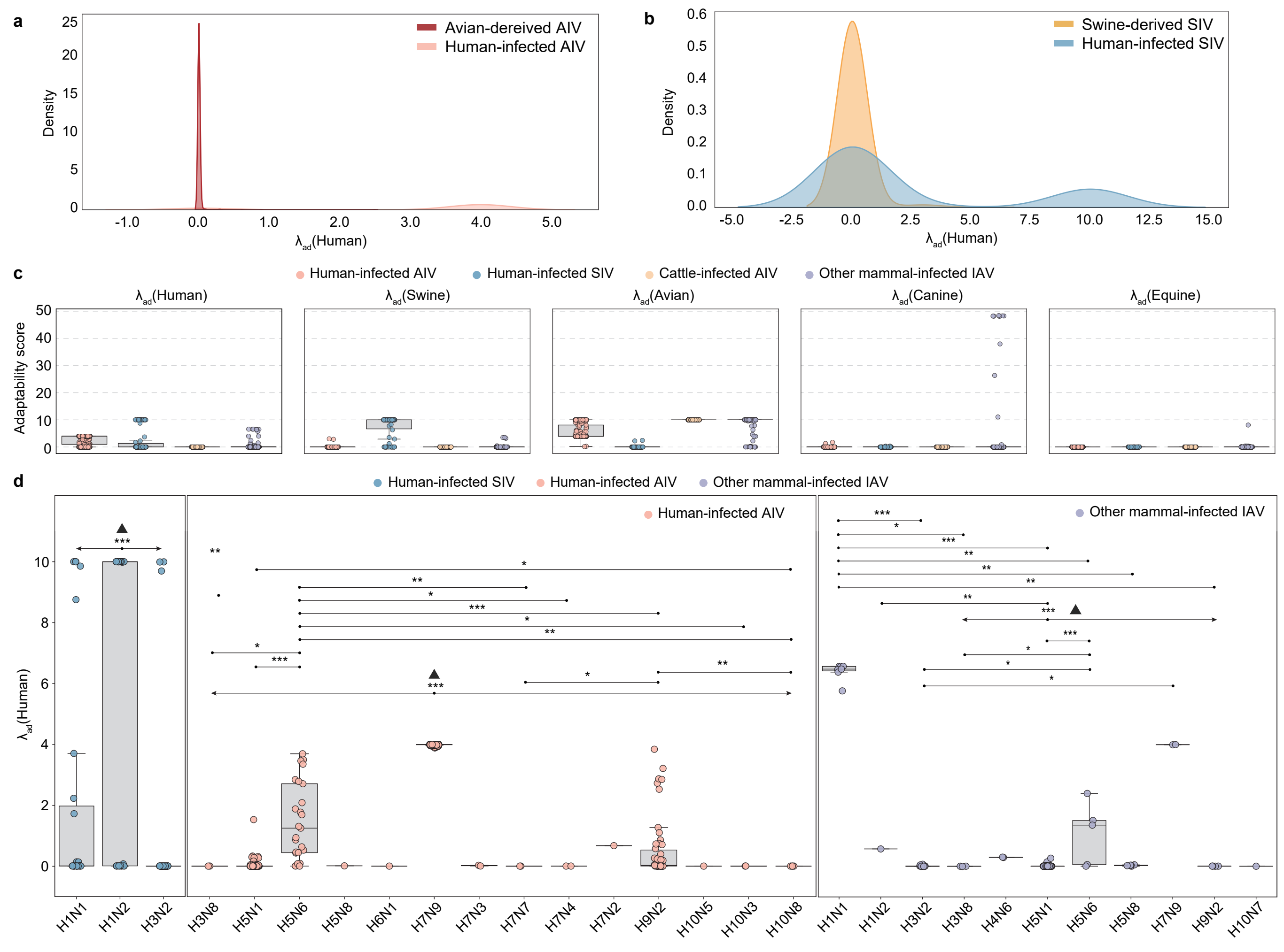

### Supplementary Fig. 13

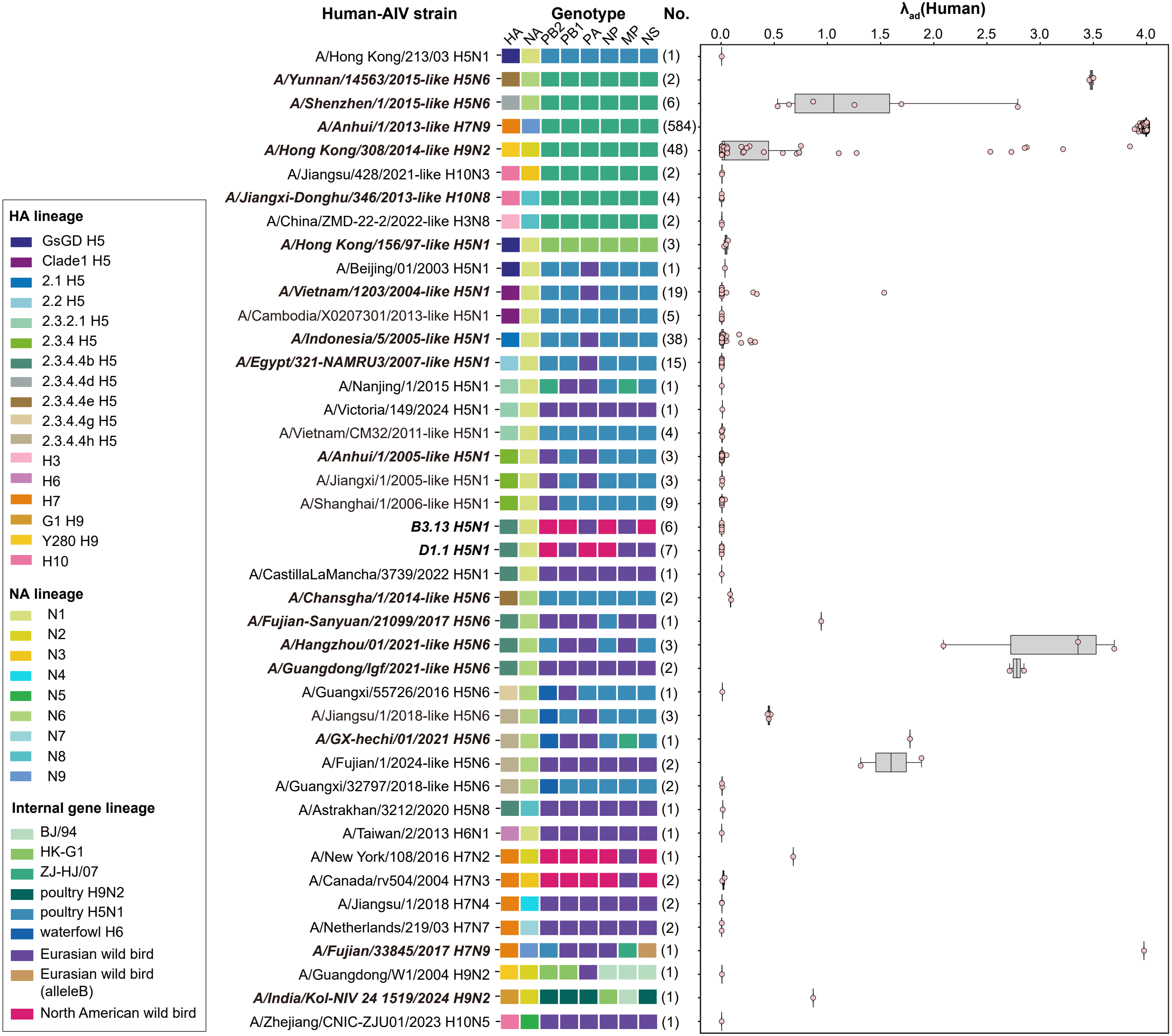

### Supplementary Fig. 14

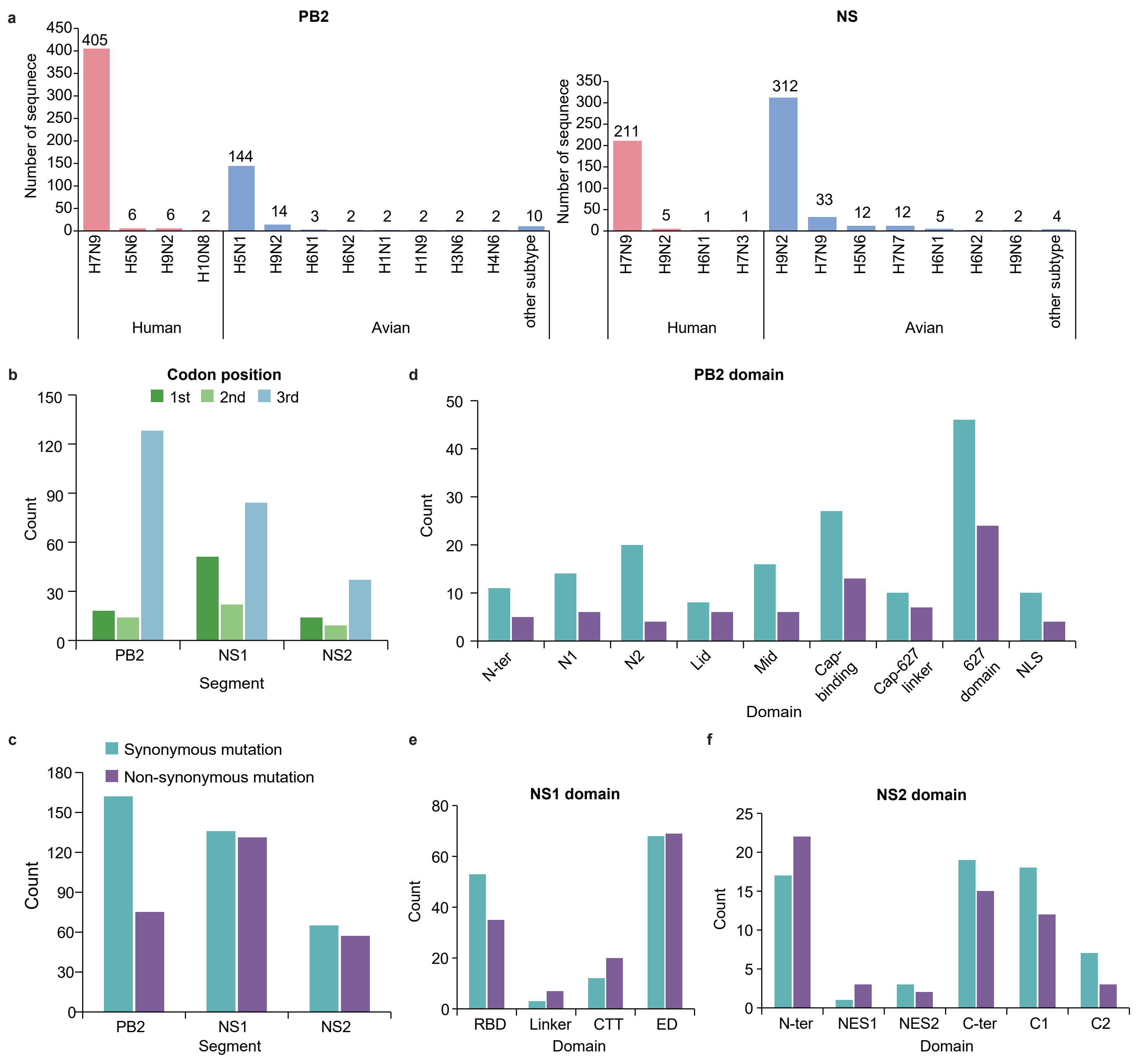
